# OTUD1 deubiquitylase regulates NF-κB- and KEAP1-mediated inflammatory responses and reactive oxygen species-associated cell death pathways

**DOI:** 10.1101/2021.12.26.474226

**Authors:** Daisuke Oikawa, Min Gi, Hidetaka Kosako, Kouhei Shimizu, Hirotaka Takahashi, Masayuki Shiota, Shuhei Hosomi, Keidai Komakura, Hideki Wanibuchi, Daisuke Tsuruta, Tatsuya Sawasaki, Fuminori Tokunaga

## Abstract

Deubiquitylating enzymes (DUBs) regulate numerous cellular functions by removing ubiquitin modifications. We examined the effects of 88 human DUBs on linear ubiquitin chain assembly complex (LUBAC)-induced NF-κB activation, and identified OTUD1 as a potent suppressor. OTUD1 regulates the canonical NF-κB pathway by hydrolysing K63-linked ubiquitin chains from NF-κB signalling factors, including LUBAC. OTUD1 negatively regulates the canonical NF-κB activation, apoptosis, and necroptosis, whereas OTUD1 upregulates the interferon (IFN) antiviral pathway. The N-terminal intrinsically disordered region of OTUD1, which contains an EGTE motif, is indispensable for KEAP1-binding and NF-κB suppression. OTUD1 is involved in the KEAP1-mediated antioxidant response and reactive oxygen species (ROS)-induced cell death, oxeiptosis. In *Otud1*^-/-^-mice, inflammation, oxidative damage, and cell death were enhanced in inflammatory bowel disease, acute hepatitis, and sepsis models. Thus, OTUD1 is a crucial regulator for the inflammatory, innate immune, and oxidative stress responses and ROS-associated cell death pathways.

---

The protein ubiquitylation system, catalyzed by ubiquitin-activating enzyme (E1), ubiquitin-conjugating enzyme (E2), and ubiquitin ligase (E3), generates monoubiquitylations and various types of polyubiquitylations *via* seven internal Lys (K) residues (*i.e.*, K6, K11, K27, K29, K33, K48, and K63) and the N-terminal methionine (M1). Moreover, hybrid- and branched-ubiquitin chains, as well as chemically-modified ubiquitins, have been recently identified. These diverse ubiquitin linkages participate in numerous cellular functions, in the so called “ubiquitin code”^1^.

Deubiquitylating enzymes (DUBs) function as erasers in the ubiquitin code^2^. Approximately 100 DUBs have been identified in the human genome and classified into seven subfamilies: ubiquitin-specific protease (USP), ubiquitin C-terminal hydrolase (UCH), ovarian tumour protease (OTU), Josephins, motif interacting with ubiquitin (MIU)-containing novel DUB (MINDY)^3^, zinc finger with UFM1-specific peptidase domain protein (ZUFSP)^4^, and JAB1/MPN/MOV34 metalloenzymes (JAMM/MPN^+^). The USP, UCH, OTU, Josephins, MINDY, and ZUFSP subfamilies are cysteine proteases, whereas the JAMM/MPN^+^ proteins are zinc metalloproteases.

The linear ubiquitin chain assembly complex (LUBAC), composed of HOIL-1L (also known as RBCK1), HOIP (RNF31), and SHARPIN, is an E3 complex that specifically generates the M1-linked linear polyubiquitin chain^5, 6^. Upon stimulation by proinflammatory cytokines, such as TNF-α, LUBAC is recruited to the TNF receptor (TNFR), and conjugates M1-ubiquitin chains to NF-κB-essential modulator (NEMO) and receptor-interacting serine/threonine-protein kinase 1 (RIP1). The M1-ubiquitin chain functions as a scaffold to form the TNFR signalling complex I and recruit the canonical IκB kinase (IKK), composed of IKKα, IKKβ, and NEMO^7^. The recruited IKK molecules are activated by a *trans*-phosphorylation mechanism, leading to the activation of NF-κB signalling. Moreover, the LUBAC activity is crucial for anti-apoptosis, and genetic defects and polymorphisms of LUBAC are associated with various disorders^6^.

At present, the K11-, K63-, and M1-ubiquitin chains are known to be involved in TNFR signalling complex I^8^, and DUBs, such as Cezanne (OTUD7B), OTULIN, CYLD, and A20, are reportedly involved in NF-κB regulation^9^. Cezanne^10^ and OTULIN^11^, OTU family DUBs, selectively hydrolyze K11- and M1-ubiquitin chains, respectively. Moreover, OTULIN and CYLD bind to the N-terminal PUB domain of HOIP, which regulates LUBAC activity^12, 13^. In contrast, A20, an OTU family DUB, does not cleave linear polyubiquitin chains, and instead it binds to the M1-ubiquitin chain through the C-terminal zinc finger 7, resulting in the suppression of NF-κB activation^14^. In addition to the homotypic linear ubiquitylations, complex ubiquitylations such as M1/K63-hybrid chain^15^ and HOIL-1L mediated ester-bound monoubiquitylation^16^ have been identified in LUBAC-mediated NF-κB activation. However, the DUBs that physiologically regulate LUBAC functions have remained elusive.

In this study, we examined the effects of 88 human DUBs on LUBAC-mediated NF-κB activation, and identified an OTU-family DUB, OTUD1 (also known as DUBA7)^17^, as the most potent down-regulator. OTUD1 reportedly regulates tumour progression^18^, cell death^19, 20^, innate immune responses^21–23^, YAP signalling^24^, autoimmune disease^25^, iron transport^26^, and inhibition of colonic inflammation^27^, whereas the regulatory mechanism of OTUD1 on the NF-κB pathway has not been revealed.

## Results

### OTUD1 is a negative regulator of canonical NF-κB signalling

To comprehensively explore the DUBs involved in LUBAC-mediated NF-κB activation, we prepared 88 human DUB cDNAs and analysed their effects by a luciferase assay in HEK293T cells (Fig. 1a). Taking the NF-κB activity induced by the expression of LUBAC alone as 100%, its co-expression with 16 DUBs, such as USP10, USP32, USP9Y, and so on, upregulated the NF-κB activity. In contrast, 34 DUBs significantly downregulated the LUBAC-induced NF-κB activation. Among them, USP2, USP13, OTULIN, OTUB1, A20, OTUD6A, OTUD2, and OTUD1 showed potent (>95%) inhibitory effects. Since OTUD6A, OTUD2, and OTUD1 had stronger inhibitory effects than those of the known LUBAC-suppressive DUBs, such as CYLD, OTULIN, and A20, we performed another NF-κB luciferase assay by expressing 10-fold higher amounts of LUBAC subunits than those in Fig. 1a. As a consequence, OTUD6A, OTUD2, and OTUD1 dose-dependently suppressed the LUBAC- and/or TNF-α-induced NF-κB activation (Fig. 1b, Extended Data Fig. 1a, 1b). Since OTUD1 showed the most potent inhibitory effect, we focused on investigating its physiological functions.

**Fig. 1.**
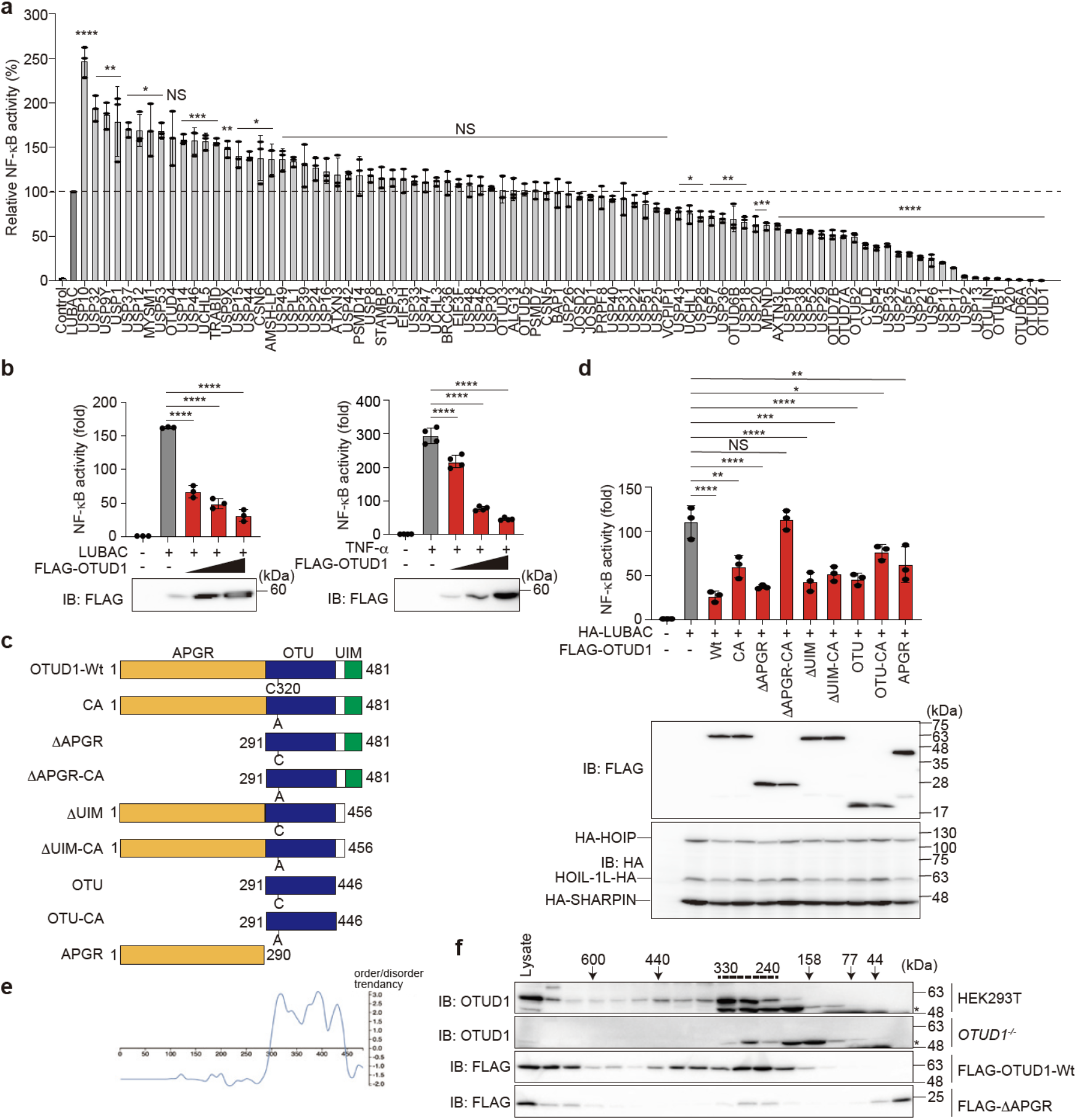
OTUD1 suppresses canonical NF-κB activation through the catalytic activity and the N-terminal region. **a,** Screening for DUBs that regulate LUBAC-mediated NF-κB activation. Effects of 88 human DUBs on LUBAC-induced NF-κB activation were analysed by a luciferase assay. **b,** Dose-dependent inhibition by OTUD1 on LUBAC- and TNF-α-induced NF-κB activation. Effects of increasing amounts (0.1, 0.3, and 1.0 μg) of OTUD1 were examined with co-expression of LUBAC or 6 h treatment with 10 ng ml^-1^ TNF-α in HEK293T cells. **c,** Domain structure of wild-type (Wt) and mutants of OTUD1. APGR: Ala-, Pro-, and Gly-rich region; OTU: ovarian tumour protease; UIM: ubiquitin-interacting motif. **d,** Effect of OTUD1 mutants on the LUBAC-induced NF-κB activity. The relative NF-κB activity induced in the presence of Wt or various mutants of OTUD1, and expression levels of OTUD1 and LUBAC subunits are shown. **a**, **b**, **d,** Data are shown as mean ± SD by ANOVA post-hoc Tukey test (*n* = 3 or 4). *: *P*<0.05, **: *P*<0.01, ***: *P*<0.001, ****: *P*<0.0001, NS: not significant. **e,** The N-terminal APGR region is disordered. Intrinsically ordered and disordered segments of OTUD1 were analysed by DICHOT^28^ (https://idp1.force.cs.is.nagoya-u.ac.jp/dichot/). **f,** OTUD1 eluted in the high molecular weight fractions. Gel filtration analyses of lysates prepared from parental and *OTUD1^-/-^* cells, and FLAG-OTUD1-Wt- and FLAG-ΔAPGR-expressing HEK293T cells were performed using a Superdex 200 column. Concentrated fractions were subjected to immunoblotting with the indicated antibodies. *: non-specific signal.

Human OTUD1 (481 a.a.) is composed of an unknown N-terminal region (a.a. 1-290), an OTU-type Cys protease domain (a.a. 291-446), which contains the active center Cys320, and a ubiquitin-interacting motif (UIM, a.a. 457-481) (Fig. 1c)^17^. Although the N-terminal region lacks homology with known domains, it shares 17% identity and 67% similarity with the amino acid sequence of the hypothetical protein Rv1157c of *Mycobacterium tuberculosis* (NP_215673.1) (Extended Data Fig. 1c), although its physiological significance remains unknown. Interestingly, the region contains an abundance of Ala (63/290 residues; 21.7%), Pro (49/290 residues; 16.9%), and Gly (28/290 residues; 9.6%). Therefore, we named the region as the Ala-, Pro-, and Gly-rich region (APGR).

To identify the critical region of OTUD1 in NF-κB inhibition, we constructed various mutants of OTUD1 and performed luciferase assays (Fig. 1c, 1d, Extended Data Fig. 1d). The OTUD1-Wt strongly suppressed the LUBAC- and TNF-α-induced NF-κB activities, and the catalytically inactive mutant of OTUD1 (CA) showed partial NF-κB suppression. The combined mutation with the catalytically inactive APGR-deletion (ΔAPGR-CA) completely abolished the NF-κB suppressive effect of OTUD1. These results suggested that both the catalytic activity and the APGR play a role in NF-κB suppression.

The protein disorder prediction by DICHOT^28^ suggested that APGR is an intrinsically disordered low complexity domain (Fig. 1e). Moreover, the endogenous and transiently expressed FLAG-tagged OTUD1 (calculated molecular weight: 51 kDa) in HEK293T cells eluted in broad fractions in a gel filtration analysis, mainly at approximately 330-240 kDa (Fig. 1f). In contrast, FLAG-ΔAPGR (calculated molecular weight: 22 kDa) predominantly eluted at monomer fractions of <44 kDa. These results suggested that the low complexity APGR functions as a multiple protein-interaction site.

### OTUD1 preferentially cleaves K63-linked ubiquitin chain

OTUD1 primarily cleaves K63-linked diubiquitin^17^, but it also cleaves K11- and K48-chains at higher concentrations^29^. The DUB protein array analysis showed that OTUD1 preferentially cleaves the K63-ubiquitin chain, followed by the K6- and K48-chains^30^. However, several cellular analyses suggested that OTUD1 cleaves K33-linked polyubiquitin chains in SMAD7^18^, and atypical K6-, K11-, and K29-linked polyubiquitin chains from IRF3 ^21^. Our biochemical analyses reconfirmed that the transiently expressed and immunoprecipitated full-length FLAG-OTUD1-Wt in HEK293T cell lysates, but not the catalytically inactive mutant (FLAG-OTUD1-CA), effectively converted K63-linked diubiquitin to monoubiquitin, among the 8 types of diubiquitins tested *in vitro* (Extended Data Fig. 1e). Moreover, recombinant GST-OTUD1 completely hydrolysed the K63-linked polyubiquitin chain, and processed a long K48-linked polyubiquitin chain to tetraubiquitin (Extended Data Fig. 1f). However, OTUD1 did not cleave di- and poly-M1 chains, indicating that OTUD1 downregulates the NF-κB activation independently of the cleavage of a LUBAC-generated linear ubiquitin chain, and principally by hydrolysing the K63-linked ubiquitin chain.

### OTUD1 down-regulates inflammatory cytokine-induced canonical NF-κB activation

The *OTUD1* gene is composed of a single exon in both human and mouse. To further clarify the physiological function of OTUD1, we constructed *OTUD1*-knockout (*OTUD1^-/-^*) HeLa and HEK293T cells and *Otud1^-/-^*-mice by the CRISPR/Cas9 technique (Extended Data Fig. 2a, 2b). The genetic depletion of *OTUD1* in HEK293T cells showed no effect on the expression of LUBAC subunits and NF-κB signalling factors, such as p105 and IκBα (Extended Data Fig. 2c). Moreover, Otud1 is ubiquitously expressed in various mouse tissues (Extended Data Fig. 2d).

To examine the cellular functions of OTUD1, we analysed the TNF-α-induced canonical NF-κB pathway. The genetic ablation of *OTUD1* in HEK293T cells enhanced the TNF-α-induced NF-κB luciferase activity (Extended Data Fig. 2e). Upon TNF-α stimulation, multiple proteins, such as RIP1, TNF receptor-associated protein with a death domain (TRADD), IKK complex, and LUBAC, are recruited by TNFR to form signalling complex I^8, 31^. In *OTUD1^-/-^*-HeLa cells, TNF-α-induced phosphorylation of IκBα, a hallmark of NF-κB activation, was upregulated as compared to that in the parental cells (Fig. 2a). Moreover, the FLAG-TNF-α precipitation indicated that polyubiquitylation and RIP1 and LUBAC recruitment were enhanced in *OTUD1^-/-^*-cells, suggesting that OTUD1 is involved in the regulation of TNFR complex I (Fig. 2a). TANK-binding kinase 1 (TBK1) and IKKε reportedly prevent TNF-induced cell death by RIP1 phosphorylation^32^. Importantly, the recruitment and phosphorylation of TBK1 and IKKε were reduced in the TNFR signalling complex I from *OTUD1^-/-^* cells (Extended Data Fig. 2f). Thus, OTUD1 reciprocally regulates TNF-α-mediated NF-κB activation and TBK1/IKKε-mediated signalling in HeLa cells. We further investigated the effect of the genetic ablation of *OTUD1* on TNF-α-induced K63-ubiquitylation, by using a tandem ubiquitin binding entity (TUBE)^33^. After TNF-α stimulation, the K63-polyubiquitylations of NEMO and RIP1 were enhanced in *OTUD1^-/-^*-HeLa cells (Fig. 2b), indicating that OTUD1 regulates K63-ubiquitylation in complex I. Indeed, the expression of NF-κB target genes was enhanced in TNF-α-treated *OTUD1*^-/-^-HeLa cells (Extended Data Fig. 2g).

**Fig. 2.**
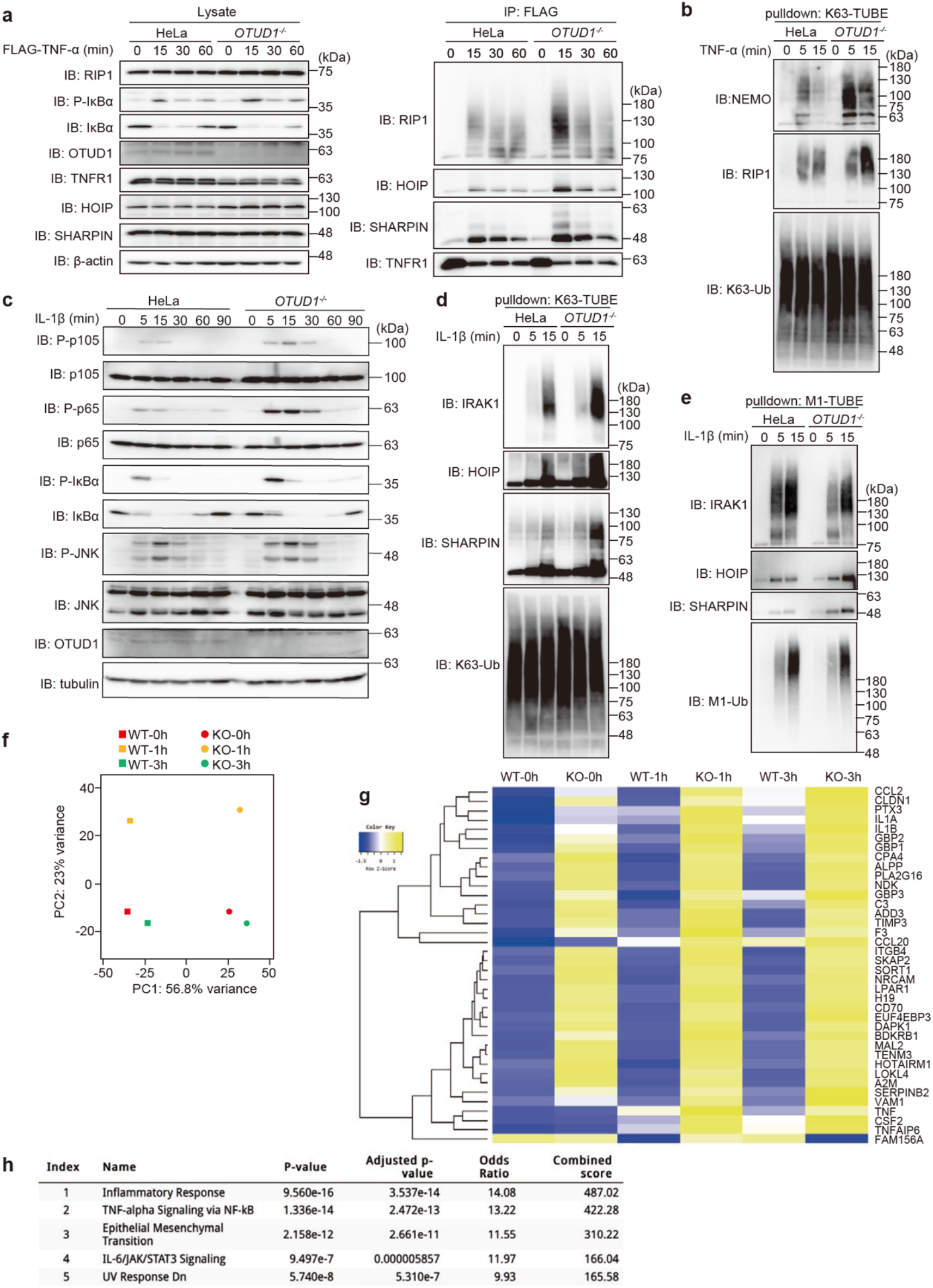
OTUD1 downregulates inflammatory cytokine-induced canonical NF-κB activation. **a,** The enhanced ubiquitylation and recruitment of NF-κB signalling factors to TNFR complex I. Parental and *OTUD1*^-/-^-HeLa cells were stimulated with 1 μg ml^-1^ FLAG-TNF-α for the indicated periods, and cell lysates and anti-FLAG immunoprecipitates were subjected to immunoblotting with the depicted antibodies. **b,** K63-linked polyubiquitylation of NF-κB activators is enhanced in *OTUD1*^-/-^ cells. Parental and *OTUD1*^-/-^-HeLa cells were stimulated with 20 ng ml^-1^ TNF-α for the indicated periods, and then pulled down by K63-TUBE. Samples were subjected to immunoblotting with the indicated antibodies. **c,** The enhanced IL-1β-induced NF-κB activation in *OTUD1*^-/-^ cells. Parental and *OTUD1*^-/-^ cells were stimulated with 1 ng ml^-1^ IL-1β, and analysed as in **a**. **d, e,** Alterations of K63- and M1-linked ubiquitylation of NF-κB activators in *OTUD1*^-/-^-HeLa cells. Parental and *OTUD1*^-/-^ cells were stimulated with 1 ng ml^-1^ IL-1β for the indicated periods, and pulled down by K63-TUBE (**d**) or M1-TUBE (**e**). Samples were immunoblotted with the indicated antibodies. **f, g,** RNA-seq analysis. Parental and *OTUD1^-/-^*-HeLa cells were stimulated with 10 ng ml^-1^ IL-1β for 1 and 3 h. The cells were lysed, and subjected to a transcriptome-wide expression analysis using their extracted total RNA. The principal component analysis (**f**) and heatmap analysis of significantly varied 34 mRNAs, using the cut off value of 10 (**g**) were performed. **h**, The enhanced inflammatory responses in *OTUD1^-/-^*-HeLa cells. Taking the cut off value of 3.0, pathway analyses of 175 up-regulated genes were performed by MSigDB Hallmark 2020, and are listed in ascending order of *P*-value.

Similarly, upon stimulation with IL-1β, the phosphorylation of NF-κB factors, such as p105, p65, and IκBα, and the subsequent degradation of IκBα, were enhanced in *OTUD1^-/-^* cells as compared to the parental cells (Fig. 2c). In contrast, the phosphorylation of JNK, a MAP kinase, was not affected in *OTUD1^-/-^*-cells. When K63-polyubiquitinated proteins were captured from cell lysates by K63-TUBE, enhanced high molecular weight smear migrations of interleukin-1 receptor-associated kinase 1 (IRAK1), HOIP, and SHARPIN were detected in IL-1β-treated *OTUD1^-/-^*-HeLa cells (Fig. 2d). In contrast, the M1-TUBE analysis indicated that the linear ubiquitylation of these factors was not increased (Fig. 2e), suggesting that OTUD1 physiologically regulates the IL-1β-induced K63-deubiquitylation of IRAK1 and LUBAC. After IL-1β-treatment, the mRNA (*IL-6*, *ICAM1*, and *BIRC3*) and protein (VCAM and IκBζ) levels of NF-κB targets were enhanced in *OTUD1^-/-^*-cells, as compared to those in parental cells (Extended Data Fig. 2h, 2i). Interestingly, the expression level of OTUD1 gradually increased after IL-1β stimulation, suggesting that OTUD1 is an NF-κB target gene.

To further investigate the effect of OTUD1 on NF-κB signalling, we performed an RNA sequencing (RNA-seq) analysis, and compared the transcripts in IL-1β-treated parental and *OTUD1^-/-^*-HeLa cells. A principal component analysis clearly demonstrated that the *OTUD1*-deficiency affected the transcription of a set of genes (Fig. 2f). The heatmap, molecular signature database (MSigDB), and KEGG pathway analyses revealed the significant upregulation of inflammatory responses and NF-κB-related factors in IL-1β-stimulated *OTUD1^-/-^*-HeLa cells (Fig. 2g, 2h, Extended Data Fig. 3a). Collectively, these results indicated that OTUD1 is a negative regulator for the IL-1β-induced canonical NF-κB activation pathway.

We further examined the role of Otud1 using splenic B cells from Wt- and *Otud1^-/-^*-mice, and confirmed that the *Otud1*-deficiency did not affect the CD40-or B cell receptor-mediated induction of NF-κB target genes, such as *Tnfaip3* (A20) and *Bcl-xl* (Extended Data Fig. 3b, 3c). Furthermore, lymphotoxin β-mediated non-canonical NF-κB activation, which is demonstrated by the intranuclear translocation of p52 and RelB, was not affected in *Otud1^-/-^*-MEFs (Extended Data Fig. 3d). Thus, Otud1 does not play a crucial role in B cells and the non-canonical NF-κB activation pathway.

### Otud1 activates IFN antiviral signalling

To further investigate the effect of Otud1 on innate immune responses, we examined the antiviral pathway using *Otud1^+/+^*- and *Otud1^-/-^*-MEFs. Upon stimulation with poly(dA:dT), which activates the IFN antiviral pathway^34^, the phosphorylation of TBK1 and IRF3 and the expression of IRF3-target genes, such as *Isg15* and *Isg56*, were reduced in *Otud1^-/-^*-MEF cells as compared to the *Otud1^+/+^*-MEFs (Fig. 3a, 3b). These results indicated that in contrast to NF-κB signalling, Otud1 functions as a positive-regulator in the IFN antiviral pathway. Similarly, the TLR3 ligand poly(I:C)-induced phosphorylation of IRF3 and the expression of IRF3-target genes were decreased in *Otud1^-/-^*-bone marrow derived macrophages (BMDMs) and MEFs (Fig. 3c, 3d, Extended Data Fig. 4a). Upon infection with Sendai virus (SeV), a single-stranded RNA virus, *Otud1^-/-^*-MEFs and -BMDMs showed reduced expression of IRF3-target genes, such as *interferon β* (*Ifnb*), *Isg15, Isg56*, and *Cxcl10,* as compared to *Otud1^+/+^*-cells (Fig. 3e, 3f). Indeed, an RNA-seq analysis of poly(I:C)-treated *Otud1^+/+^*- and *Otud1^-/-^*-MEF cells revealed that the *Otud1*-deficiency downregulated the transcription of a set of IFN genes (Fig. 3g, 3h). MSigDB and KEGG pathway analyses demonstrated that the *Otud1*-deficiency affected the expression of IFN and the RIG-I-like receptor signalling pathway after poly(I:C) stimulation (Fig. 3i, Extended Data Fig. 4b). Collectively, these results indicated that Otud1 is a positive regulator for the type I IFN antiviral pathway.

**Fig. 3.**
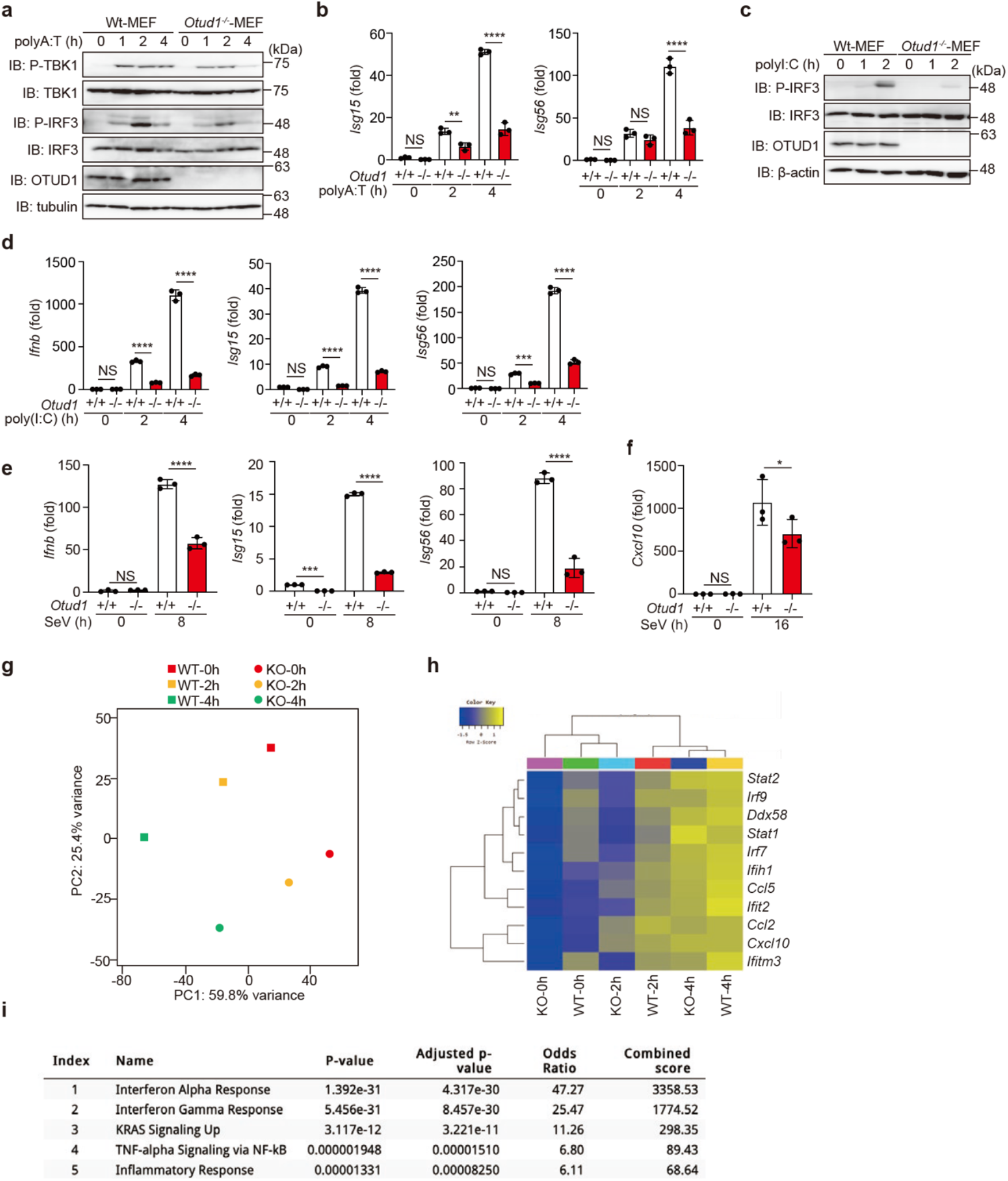
OTUD1 upregulates type I IFN antiviral pathway. **a,** The poly(A:T)-mediated IFN antiviral pathway is suppressed in *Otud1^-/-^*-MEFs. *Otud1^+/+^*- and *Otud1^-/-^*-MEF cells were stimulated with 1 μg ml^-1^ poly(A:T) for the indicated periods, and the cell lysates were immunoblotted with the indicated antibodies. **b,** The expression of IRF3-target genes is suppressed in poly(A:T)-treated *Otud1^-/-^*-MEF cells. MEFs were stimulated with 1 μg ml^-1^ poly(A:T) for the indicated periods, and the mRNA levels were assessed by qPCR. **c,** The suppressed poly(I:C)-mediated antiviral response in *Otud1^-/-^*-MEFs. *Otud1^+/+-^*- and *Otud1^-/-^*-MEFs were stimulated with 10 μg ml^-1^ poly(I:C) for the indicated periods, and cell lysates were immunoblotted with the indicated antibodies. **d,** Reduced poly(I:C)-mediated antiviral response in *Otud1^-/-^*-BMDMs. BMDMs from *Otud1^+/+-^*- and *Otud1^-/-^*-mice were stimulated with 10 μg ml^-1^ poly(I:C) for the indicated periods, and a qPCR analysis was performed. **e, f**, Sendai virus (SeV)-induced antiviral response is suppressed in *Otud1^-/-^*-cells. *Otud1^+/+-^*- and *Otud1^-/-^*-MEFs (**e**) or BMDMs (**f**) were infected with SeV at a CIU of 10 for 1 h, incubated for the indicated periods, and then subjected to qPCR analyses. **b**, **d**, **e**, **f**, Data are shown as mean ± SD by ANOVA post-hoc Tukey test (*n* = 3). *: *P*<0.05, **: *P*<0.01, ***: *P*<0.001, ****: *P*<0.0001, NS: not significant. **g-i** Suppressed transcription of IRF3-target genes in *Otud1^-/-^*-MEFs. *Otud1^+/+^*- and *Otud1^-/-^*-MEFs were stimulated with 10 μg ml^-1^ poly(I:C) for 2 and 4 h, and RNA-seq analyses were performed. **g,** A principal component analysis of the RNA-seq analysis. **h**, Taking the cutoff value of 10, significantly varied mRNAs are shown by the heatmaps. **i**, Taking the cutoff value of 3.0, pathway analyses of 179 down-regulated genes were performed by MSigDB Hallmark 2020, and listed in ascending order of *P*-value.

### OTUD1 suppresses TNF-α-induced apoptosis and necroptosis

When TNFR signalling complex I fails to induce NF-κB-mediated cell survival under conditions where protein synthesis is inhibited by cycloheximide, TNF-α stimulation induces cell death with the formation of a second complex, named complex II, which is composed of Fas-associated death domain protein (FADD) and caspase 8^35, 36^. A pan-caspase inhibitor (ZVAD) suppresses apoptosis, whereas it induces another programmed necrosis, necroptosis. The kinase activity of RIP1 subsequentially phosphorylates RIP3 and then mixed lineage kinase domain-like pseudokinase (MLKL), which is indispensable for necroptosis^37^, and therefore, the RIP1 kinase inhibitor, necrostatin-1 (Nec-1), suppresses necroptosis^38^.

After an 8 h treatment with TNF-α+cycloheximide (TC), the trypan blue-positive dead cells were significantly increased in *OTUD1^-/-^*-HeLa and HEK293T cells, which both lack RIP3 expression (Fig. 4a). The cell survival analysis by an xCELLigence real-time cell analyser indicated that *OTUD1^-/-^*-HEK293T cells are more sensitive to cell death than the parental cells after TC-treatment (Fig. 4b). Indeed, the cleavages of PARP, caspase 3, and caspase 8, hallmarks of the extrinsic apoptotic pathway, were enhanced in TC-treated *OTUD1^-/-^*-HEK293T cells as compared to those in parental cells (Fig. 4c, 4d). When cell lysates were immunoprecipitated by an anti-FADD antibody, effective complex II formation, as shown by the enhanced association of caspase 8 with FADD, was detected in *OTUD1^-/-^*-HEK293T cells (Fig. 4d). These results suggested that OTUD1 has an inhibitory effect on the TNF-α-induced apoptosis.

**Fig. 4.**
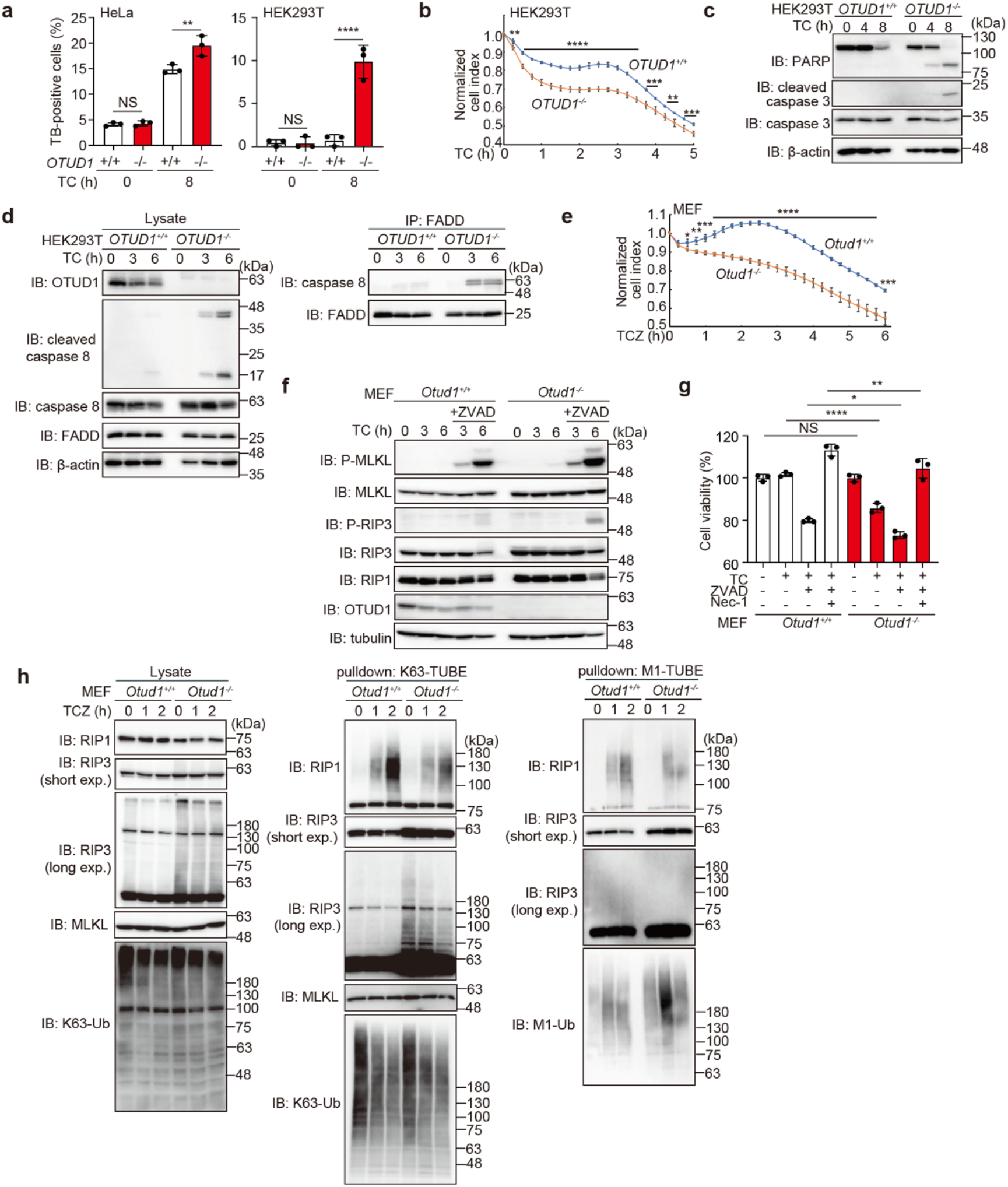
OTUD1 regulates TNF-α-induced apoptosis and necroptosis. **a,** Increased TNF-α-induced cell death by the genetic ablation of *OTUD1.* Wt- and *OTUD1*^-/-^-HeLa and HEK293T cells were treated with or without 10 ng ml^-1^ TNF-α and 10 μg ml^-1^ CHX (TC), and trypan blue-positive cells were counted. **b,** Reduced cell viability of *OTUD1*-deficient cells. Wt- and *OTUD1*^-/-^-HEK293T cells were treated with TC, and the cell proliferation was measured by xCELLigence real-time cell monitoring. **c,** Enhanced apoptosis in *OTUD1*^-/-^-HEK293T cells. Wt- and *OTUD1*^-/-^-HEK293T cells were treated with TC, and cell lysates were immunoblotted with the indicated antibodies. **d,** TNFR complex II formation was accelerated in *OTUD1^-/-^*-HEK293T cells. Wt- and *OTUD1^-/-^*-HEK293T cells were treated with TC as in **c**. The cell lysates and anti-FADD immunoprecipitates were then blotted with the indicated antibodies. **e,** Increased necroptosis in *Otud1^-/-^*-MEFs*. Otud1^+/+^*- and *Otud1^-/-^*-MEFs were treated with 10 ng ml^-1^ TNF-α, 10 μg ml^-1^ CHX, and 20 μM ZVAD (TCZ), and the cell proliferation was measured as in **b**. **f,** Increased phosphorylation of MLKL and RIP3 in TCZ-treated *Otud1^-/-^*-MEFs. MEFs were treated with TC with or without ZVAD as indicated, and the cell lysates were immunoblotted with the depicted antibodies. **g,** Necrostatin-1 (Nec-1) rescued cell death. *Otud1^+/+^*- and *Otud1^-/-^*-MEFs were treated with TC, ZVAD, and/or 100 μM Nec-1 for 8 h, as indicated, and cell viability was analysed by a Celltiter Glo assay. **h,** Enhanced K63-ubiquitylation of RIP3 in *Otud1^-/-^*-MEFs. MEFs were treated with TCZ for the indicated periods, and cell lysates and K63-or M1-TUBE precipitates were immunoblotted with the indicated antibodies. Data are shown as mean ± SD by ANOVA post-hoc Tukey test (**a**, **g**; *n* = 3) or *t*-test at each time point (**b**, **e**; n = 4). *: *P*<0.05, **: *P*<0.01, ***: *P*<0.001, ****: *P*<0.0001, NS: not significant.

We next investigated the involvement of Otud1 in necroptosis, using MEFs. In the presence of TC with ZVAD (TCZ), the *Otud1^-/-^*-MEFs died more rapidly than the *Otud1^+/+^*-MEFs (Fig. 4e). The phosphorylation of MLKL and RIP3 was enhanced in *Otud1^-/-^*-MEFs (Fig. 4f), and TCZ-induced cell death in *Otud1^-/-^*-MEFs was rescued by Nec-1 (Fig. 4g). Interestingly, the polyubiquitylation of RIP3, but not RIP1, was enhanced in *Otud1^-/-^*-MEFs (Fig. 4h, left panel), indicating that RIP3 is a possible endogenous substrate of Otud1. When K63-ubiquitinated proteins were pulled down by K63-TUBE, the smeared migration of RIP1 was slightly suppressed upon stimulation with TCZ in *Otud1^-/-^*-MEFs (Fig. 4h, middle panel). In contrast, the M1-ubiquitylation of RIP3 was not detected in either MEFs (Fig. 4h, right panel). These results indicated that Otud1 is a critical regulator in TNF-α-mediated necroptosis pathways through the removal of K63-ubiquitin chains from RIP3 under steady state conditions.

### OTUD1 regulates KEAP1-mediated oxidative stress response and cell death

To further examine the physiological roles of OTUD1, proteins interacting with FLAG-OTUD1 in HEK293T cells were analysed by immunoprecipitation-mass spectrometry. We identified KEAP1 as the most prominent interactor of OTUD1 (Fig. 5a, Extended Data Fig. 5a). KEAP1 regulates the proteasomal degradation of the NRF2 transcription factor, as a component of the Cullin 3 (CUL3)-RING E3 complex^39^. Moreover, KEAP1 is involved in excess reactive oxygen species (ROS)-induced cell death, named oxeiptosis, with the apoptosis-inducing factor mitochondria associated 1 (AIFM1) and the mitochondrial Ser/Thr protein phosphatase PGAM5^40^. Indeed, peptides derived from AIFM1, CUL3, and PGAM5 were identified in FLAG-OTUD1 co-precipitates (Fig. 5a, Extended Data Fig. 5a). We detected the endogenous association of KEAP1 and OTUD1 (Fig. 5b), and the K63-polyubiquitylation of KEAP1 was enhanced in *Otud1^-/-^*-MEFs (Fig. 5c). Moreover, we determined that the N-terminal APGR of OTUD1 interacts with the C-terminal Kelch and CTR of KEAP1 (Fig. 5d, 5e). Importantly, we found that the human and mouse OTUD1 proteins contain ETGE and the similar DTGE sequence, respectively, in APGR, which plays a crucial role in NRF2 for KEAP1-binding^41^ (Fig. 5f). Indeed, the deletion of the ETGE motif from OTUD1 abolished the KEAP1-binding ability (Fig. 5g). Upon H2O2-treatment, ROS production was upregulated in *Otud1^-/-^*-MEFs as compared to *Otud1^+/+^*-MEFs (Fig. 5h), resulting in the enhanced expression of antioxidant genes, such as *Hmox-1* (Ho-1) and *Nhe2l2* (Nrf2) at the early phase (∼3 h) (Fig. 5i). However, a prolonged treatment with H_2_O_2_ caused drastic cell death in *Otud1^-/-^*-MEFs, suggesting that the excessive ROS production subsequently induces oxeiptosis (Fig. 5j).

**Fig. 5.**
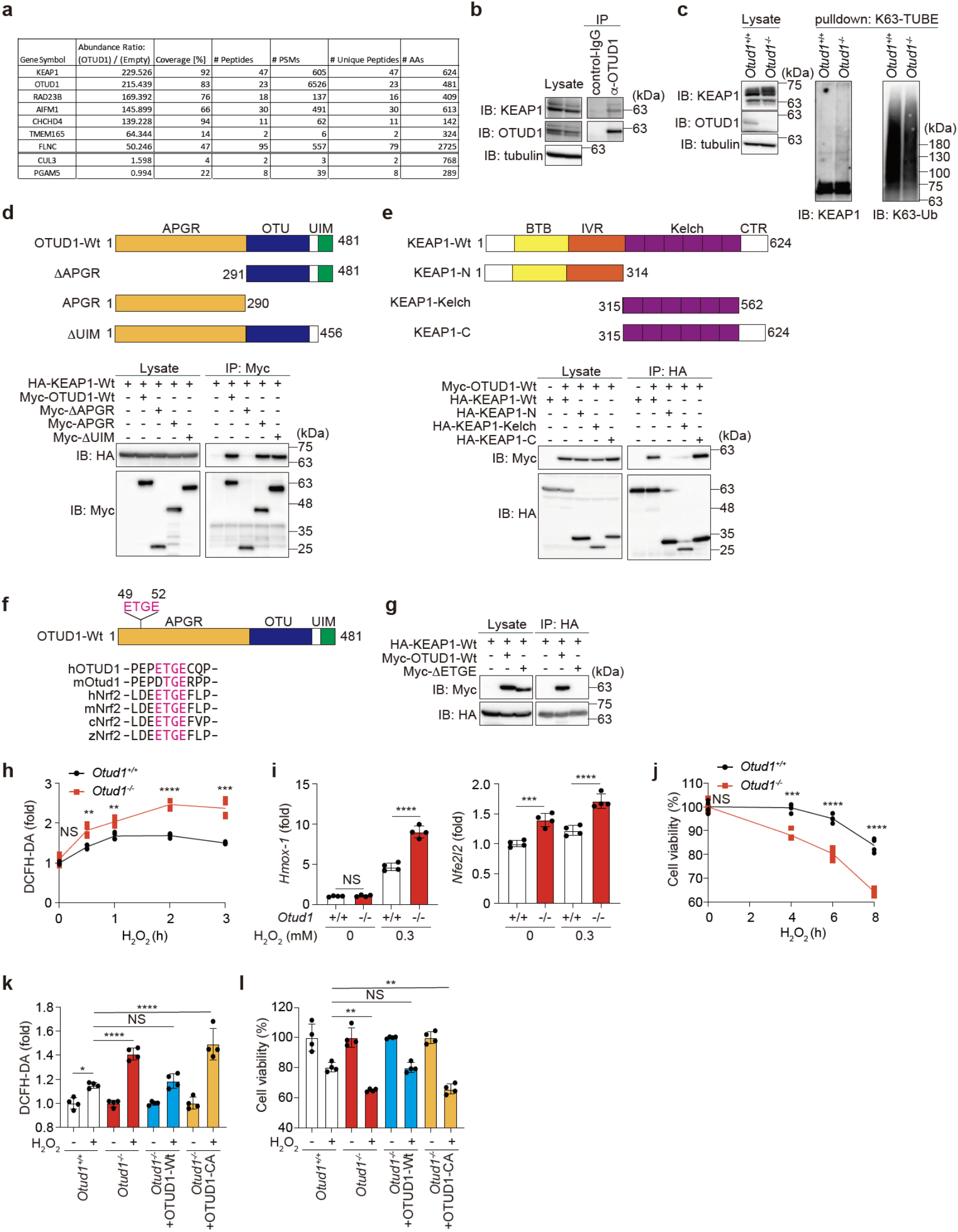
OTUD1 regulates KEAP1-mediated oxidative stress response and ROS-associated cell death. **a,** Identification of OTUD1-interacting proteins. FLAG-tagged OTUD1 was expressed in HEK293T cells and immunoprecipitated with an anti-FLAG antibody, and the co-immunoprecipitants were then analysed by MS. Proteins with an abundance ratio greater than 50, and values for CUL3 and PGAM5 are shown. **b,** OTUD1 physiologically interacts with KEAP1. HEK293T cell lysates and anti-OTUD1 immunoprecipitates were subjected to immunoblotting with the indicated antibodies. **c,** OTUD1 regulates K63-ubiquitylation of KEAP1. Cell lysates and K63-TUBE precipitates from *Otud1*^+/+^- and *Otud1*^-/-^-MEFs were immunoblotted with the indicated antibodies. **d,** APGR of OTUD1 is required for KEAP1-binding. Wt- and mutants of Myc-OTUD1 were expressed with HA-KEAP1 in HEK293T cells. Cell lysates and anti-Myc-immunoprecipitates were immunoblotted with the indicated antibodies. **e,** The Kelch domain and the C-terminal region (CTR) of KEAP1 are responsible for OTUD1-binding. A similar analysis as in **d** was performed using Wt- and mutants of HA-KEAP1 and Myc-OTUD1. **f,** OTUD1 contains an ETGE motif in APGR. Localization of the ETGE motif in OTUD1, and amino acid sequence alignment with NRF2 are shown ^41^. h: human, m: mouse, c: chicken, z: zebrafish. **g**, The ETGE motif in OTUD1 is indispensable for KEAP1-binding. Myc-tagged Wt-or ETGE motif-deleted mutant of OTUD1 was expressed with HA-KEAP1-Wt as indicated, and cell lysates and anti-HA immunoprecipitates were immunoblotted with the depicted antibodies. **h,** Enhanced hydrogen peroxide-induced ROS generation in *Otud1*-deficient cells. *Otud1^+/+^*- and *Otud1^-/-^*-MEFs were treated with 0.3 mM H_2_O_2_ for the indicated periods, and the intracellular ROS levels were analysed by a DCFH-DA assay. **i,** Enhanced expression of NRF2 target genes in *Otud1*^-/-^-MEFs. MEFs were treated with or without 0.3 mM H_2_O_2_ for 3 h, and a qPCR analysis was performed. **j**, Reduced cell viability in *Otud1*^-/-^-MEFs under oxidative stress. MEFs were treated with 0.3 mM H_2_O_2_ for the indicated periods, and cell viability was analysed by Celltiter Glo assay. **k**, DUB activity of OTUD1 is necessary to suppress ROS production. OTUD1-Wt or active site mutant (CA) was restored into *Otud1^-/-^*-MEFs, and ROS levels were analysed by a DCFH-DA assay after treatment with or without 0.3 mM H_2_O_2_ for 2h. **l**, DUB activity of OTUD1 protects cells from H_2_O_2_-induced death. A similar treatment as in **k** was performed with or without 0.1 mM H_2_O_2_ for 8 h, and cell viability was analysed by a Celltiter Glo assay. Data are shown as mean ± SD by *t*-test of each time point (*n* = 4, **h, j**) or ANOVA post-hoc Tukey test (*n* = 4, **i, k, l**). *: *P*<0.05, **: *P*<0.01, ***: *P*<0.001, ****: *P*<0.0001, NS: not significant.

Interestingly, the restoration of OTUD1-Wt, but not the catalytically inactive mutant (CA), rescued ROS resistance and cell viability (Fig. 5k, 5l), suggesting that the DUB activity of OTUD1 is necessary to resist oxeiptosis. OTUD1 directly bound with KEAP1, and PGAM5 co-precipitated with OTUD1 only in the presence of KEAP1 (Extended Data Fig. 5b). Although the co-precipitation of endogenous AIFM1 with FLAG-OTUD1 was detected by the MS analysis (Fig. 5a, Extended Data Fig. 5a), the co-precipitation of exogenous OTUD1 with AIFM1 was not detected in the presence of KEAP1 and PGAM5 (Extended Data Fig. 5b). Whereas, in *Otud1^-/-^*-MEFs, increased intranuclear AIFM1 was detected concomitantly with the reduced mitochondrial AIFM1 (Extended Data Fig. 5c), which may affect mitochondrial oxidative phosphorylation and ROS generation^20^. These results indicated that OTUD1 physiologically binds and regulates the K63-ubiquitylation of KEAP1 under basal conditions and is important for the efficient antioxidative stress response, ROS production, and oxeiptosis.

In addition to H_2_O_2_-treatment, the ROS levels were also upregulated in *Otud1^-/-^*-MEFs upon apoptotic (TC) and necroptotic (TCZ) stimuli, although TNF-α alone showed no effects on ROS production (Extended Data Fig. 5d). Since the enhanced ROS production was cancelled by the restoration of Otud1-Wt, but not -CA, in TC-or TCZ-treated *Otud1^-/-^*-MEFs, the catalytic activity of Otud1 is necessary to regulate TNF-α-induced cell death pathways (Extended Data Fig. 5e). Collectively, these results indicated that OTUD1 activity regulates ROS-associated cell death pathways, such as TNF-α-induced apoptosis and necroptosis, as well as KEAP1-mediated oxeiptosis.

### OTUD1 suppresses inflammatory and oxidative stress responses *in vivo*

A recent report showed that OTUD1 inhibits colonic inflammation *in vivo*^27^. We also confirmed that significant shortening of the colon, and higher diarrhea and fecal blood scores were observed in a DSS-administrated ulcerative colitis-like inflammatory bowel disease (IBD) model of *Otud1^-/-^*-mice (Extended Data Fig. 6a, 6b). The middle and distal colon regions in DSS-treated *Otud1^-/-^*-mice showed more severe damage, characterized by multiple focal dropouts of entire crypts, inflammatory cell infiltration, and edema, as compared with the DSS-treated *Otud1^+/+^* mice (Fig. 6a). Histologic scores of colitis were significantly higher in the middle and distal colon of DSS-treated *Otud1^-/-^*-mice, as compared with DSS-treated *Otud1^+/+^* mice (Fig. 6b). Moreover, the expressions of NF-κB target genes such as *Tnf* and *Il1b* were upregulated in the distal colon from *Otud1^-/-^*-mice (Fig. 6c). Importantly, we detected the increased oxidative DNA damage and cell death in the middle and distal colon of DSS-treated *Otud1^-/-^*-mice, as evidenced by increased numbers of 8-OHdG-positive cells and TUNEL-positive cells in the colon epithelium, respectively (Fig. 6d, 6e). In addition to the IBD-like phenotype, DSS-treated *Otud1^-/-^*-mice also showed splenomegaly with increased TUNEL-positive splenocytes (Extended Data Fig. 6c, 6d). These results indicated that Otud1 suppresses the inflammatory and oxidative damage responses in an *in vivo* colitis model.

**Fig. 6.**
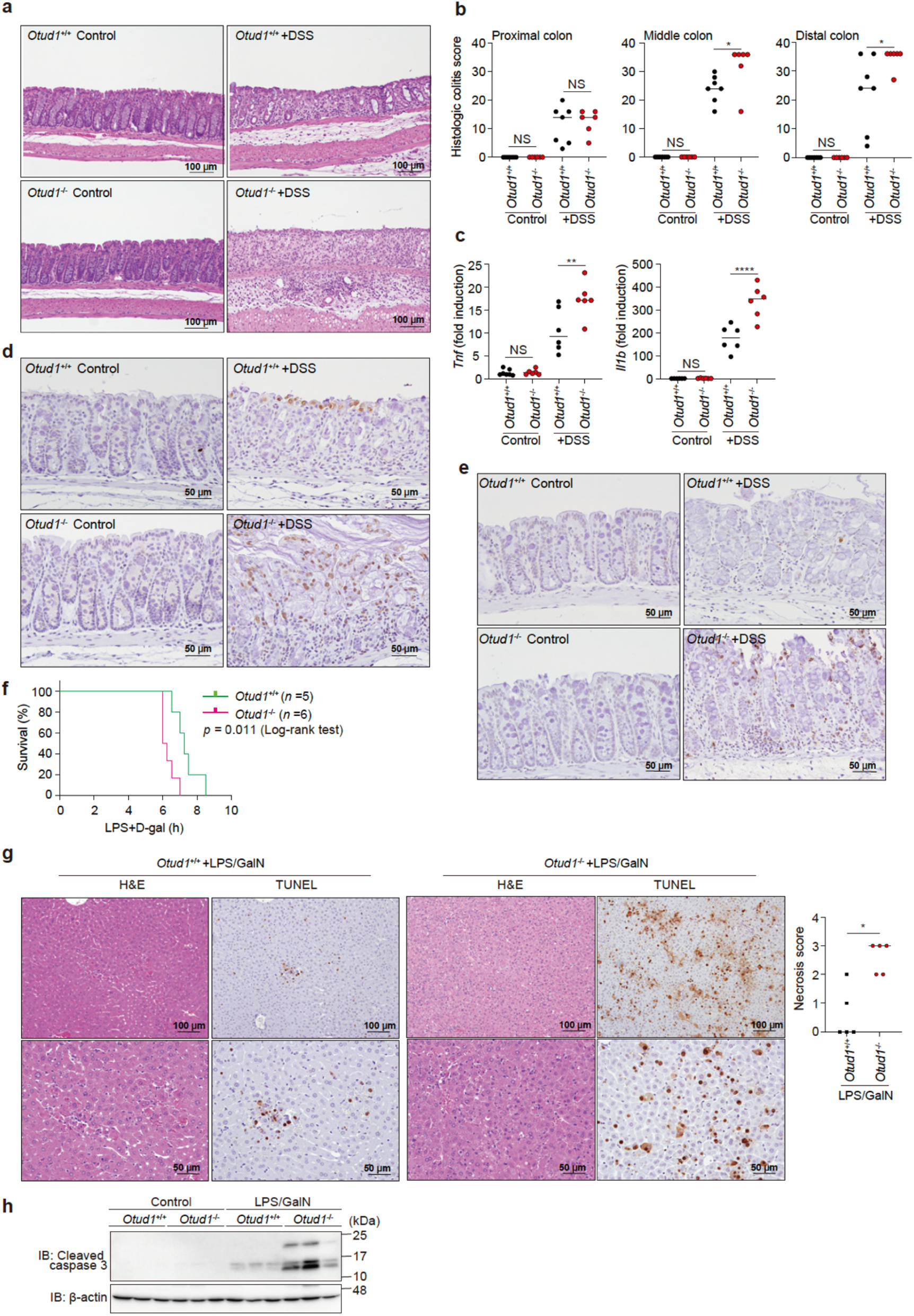
Enhanced inflammation, oxidative damage, and cell death in mouse inflammatory disease models. **a, b**, Colon damage in DSS-administered *Otud1^-/-^*-mice. *Otud1^+/+^*- and *Otud1^-/-^*-mice were treated with 2.5% DSS for 7 days, and then sacrificed. Representative images of H&E staining of distal colon (**a**) and histologic scores of colitis (**b**) (*n* = 6-7) are indicated. *Bars*; 100 µm. **c**, Increased expression of NF-κB target genes in DSS-treated *Otud1^-/-^*-mice. qPCR analysis of NF-κB targets in distal colon from DSS-treated mice (*n* = 6). **d, e**, Increased oxidative DNA damage and cell death in DSS-administered *Otud1^-/-^*-mice. Specimens as in **a** were stained for 8-OHdG (**d**) or TUNEL (**e**). *Bars*: 50 µm. **f**, *Otud1^-/-^*-mice were labile for the LPS/GalN-induced acute hepatitis model. After injections of LPS (10 μg kg^-1^) and GalN (800 mg kg^-1^), the % survivals of *Otud1^+/+^*-(*n* = 5) and *Otud1^-/-^*-(*n* = 6) mice were determined by the Kaplan-Meier method with the Log-rank test. **g,** Enhanced liver damage in *Otud1^-/-^*-mice. H&E and TUNEL staining of livers from *Otud1^+/+^*- and *Otud1^-/-^*-mice after 5 h administration of LPS/GalN, and hepatocellular death scores (*n* = 5) are shown. *Bars*: 100 µm (*upper panels*) and 50 µm (*lower panels*). **h,** Increased caspase 3 activation in hepatocytes from *Otud1^-/-^*-mice. After a 5 h treatment with LPS/GalN, livers were excised, lysed, and immunoblotted with the indicated antibodies. **b**, **c**, **g**, Data are shown as scatter plots and evaluated by Mann-Whitney test. *: *P*<0.05, **: *P*<0.01, ****: *P*<0.0001, NS: not significant.

To further clarify the pathophysiological role of Otud1, we adopted an LPS/GalN-induced acute hepatitis model. The LPS/GalN-treated *Otud1^-/-^*-mice showed significantly shorter survival, remarkably increased hepatocellular death, and increased caspase 3 activation (Fig. 6f, 6g, 6h), indicating that Otud1 protects against acute hepatitis.

In contrast to LPS/GalN, LPS challenge alone induces systemic inflammation and sepsis. When mice were intraperitoneally administered 20 mg kg^-1^ LPS, we found that the survival times of *Otud1^-/-^*-mice were significantly shorter as compared to those of *Otud1^+/+^*-mice, indicating that *Otud1^-/-^*-mice were sensitive to LPS-induced systemic inflammation and sepsis (Extended Data Fig. 6e). Histopathological findings in the livers are shown in Extended Data Table 1 and Extended Data Fig. 6f. In male mice, mild fatty changes characterized by increased microvacuoles in the cytoplasms of hepatocytes were observed in all LPS-treated *Otud1^+/+^*- and *Otud1^-/-^*-mice, but not in the *Otud1^+/+^* and *Otud1^-/-^* controls (Extended Data Fig. 6f, upper panels). Hydropic degeneration in midzonal and centrilobular hepatocytes was observed in 2 of the 6 LPS-treated male *Otud1^+/+^*-mice (Extended Data Table 1), but not in the *Otud1^+/+^* controls. No hydropic degeneration was observed in the male *Otud1^-/-^*-mice regardless of LPS treatment. Notably, in the female mice, the incidence of hepatocellular death (necrosis and/or apoptosis), predominantly localized in the midzonal area, was increased in the LPS-treated *Otud1^-/-^*-mice (5/6, 83.3%) as compared with the LPS-treated *Otud1^+/+^*-mice (2/6, 33.3%), albeit without statistical significance (Extended Data Table 1, Extended Data Fig. 6f, lower panels). This suggested that the female *Otud1^-/-^*-mice exhibited higher susceptibility to LPS-induced hepatotoxicity than their wild-type counterparts. The incidence of hydropic degeneration of hepatocytes was comparable between the LPS-treated *Otud1^+/+^* (4/6, 66.7%) and *Otud1^-/-^*-mice (3/6, 50%). Inflammatory cell infiltration was also observed in all LPS-treated mice, without apparent gender and genotype differences. In addition, the AST activities were significantly elevated in LPS-treated *Otud1^-/-^*-mice, as compared to the LPS-treated *Otud1^+/+^*-mice (Extended Data Fig. 6g). These results indicated that Otud1 is a crucial regulator of innate immune responses, and the genetic ablation of *Otud1* causes aberrant signal transduction resulting in septic shock.

### OTUD1 is associated with prognosis of kidney cancer

Finally, to investigate the involvement of OTUD1 in human diseases, we analysed cancer databases. We found that low levels of *OTUD1* expression are significantly associated with a poor prognosis in renal clear cell carcinoma patients (Fig. 7a). Importantly, the expressions of OTUD1 and KEAP1 were significantly correlated (*P* = 4.7 x 10^-9^, R = 0.54) in the normal kidney cortex; however, the positive correlation was lost in the tumour kidney cortex (Fig. 7b). In contrast, the expression levels of OTUD1 and NF-κB1 (p105/p50) were correlated in both the normal and tumour kidney cortexes. Thus, we concluded that the orchestration of OTUD1 with KEAP1 functions to suppress kidney cancer in humans, and therefore, OTUD1 is a critical regulator to maintain homeostasis.

**Fig. 7.**
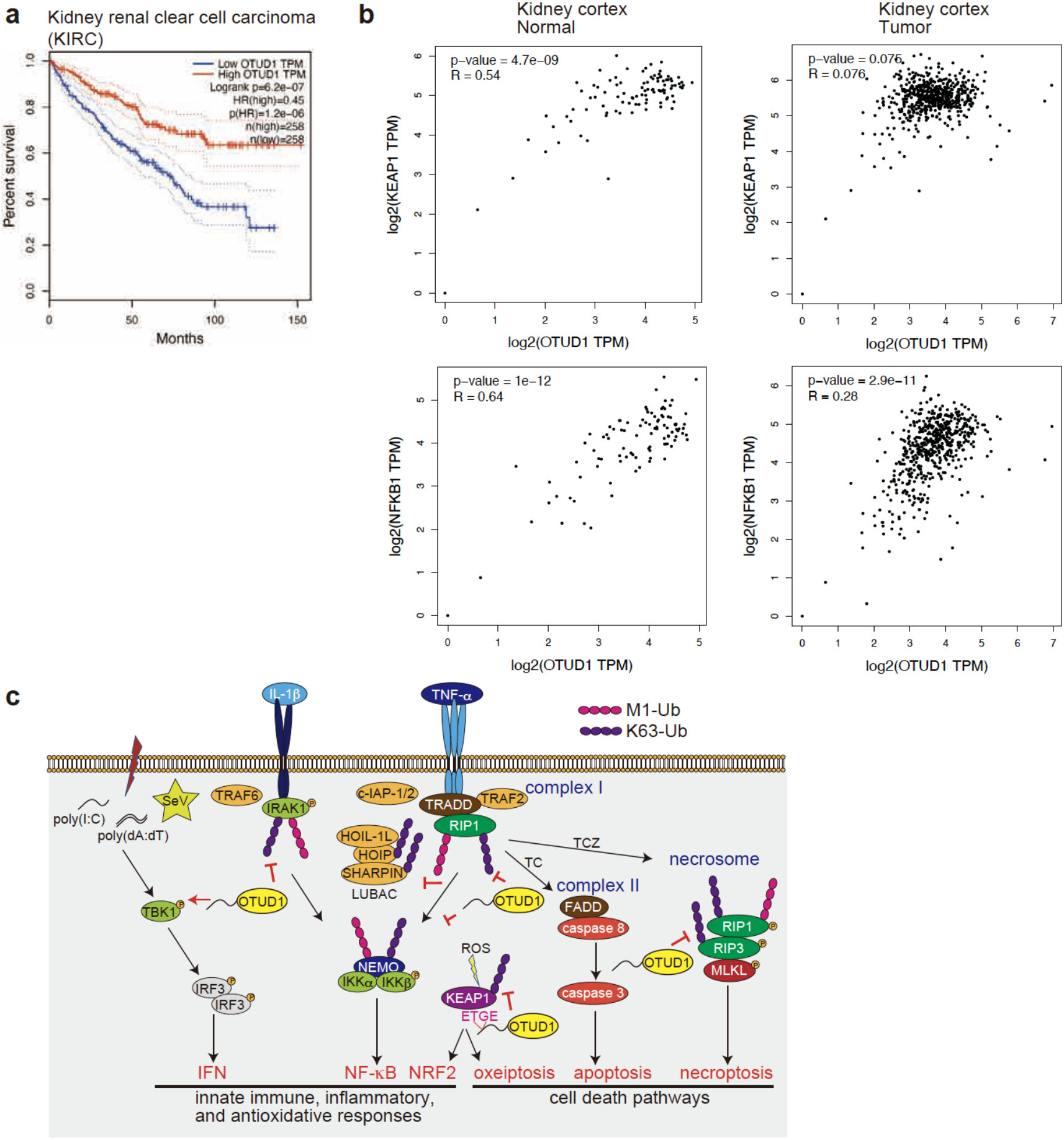
OTUD1 suppresses kidney cancer. **a**, Low expression of *OTUD1* is a poor prognosis factor in kidney cancer. Kaplan-Meier survival curves of patients with KIRC (Kidney Renal Clear Cell Carcinoma) with high (*n* = 258 patients) and low (*n* = 258 patients) expression of *OTUD1* mRNA, determined by the Log-rank test, are shown. **b**, Loss of positive correlation between *OTUD1* and *KEAP1* in kidney cancer. Correlations of expression levels with Pearson correlations and *P*-values of *OTUD1* vs*. KEAP1* (*upper panels*) or *OTUD1* vs*. NF-κB1* (*lower panels*) in normal (*left panels*) and tumour (*right panels*) kidney cortex are shown. **c**, Schematic representation of OTUD1 functions in the regulation of inflammatory and KEAP1-mediated oxidative stress responses, and cell death pathways.

## Discussion

We have identified that USP2, USP13, OTULIN, OTUB1, A20, OTUD6A, OTUD2, and OTUD1, among 88 human DUBs, exert potent inhibitory effects on LUBAC-induced NF-κB activation (Fig. 1a). Indeed, USP2 reportedly down-regulates the NF-κB pathway by removing ubiquitin chains generated by TRAF2 and TRAF6^42–44^, and the genetic ablation of *USP13* enhances the NF-κB and IFN antiviral pathways by deubiquitylating STING^45^. OTULIN and A20 are well-known down-regulators of the LUBAC-mediated NF-κB pathway^9^. OTUB1 affects NF-κB signalling by associating with TRAF3, TRAF6, and c-IAP-1^46, 47^, and OTUD2 interacts with TRAF6 during IL-1β-induced NF-κB activation^48^. Thus, most of the listed DUBs are known as regulators for the innate immune and NF-κB pathways.

OTUD1 was identified as the most potent inhibitor for LUBAC-induced NF-κB activation (Fig. 1). OTUD1 was initially found as a biomarker of thyroid cancer^49^ that reportedly deubiquitinates and stabilizes the p53 tumour suppressor^50^. OTUD1 also upregulates the expression of p21 and Mdm2, thus accelerating apoptosis^50^, and cleaves K33-linked polyubiquitin chains from the TGF-β pathway inhibitor SMAD7, which is involved in the repression of breast cancer metastasis^18^. Furthermore, OTUD1 is induced by RNA viruses, which gives rise to the upregulation of the intracellular amounts of Smurf1 through deubiquitylation ^51^. Since Smurf1 leads to the ubiquitylation-dependent degradation of MAVS, TRAF3, and TRAF6, subsequently suppressing IFN production, the genetic deficiency of *Otud1* in mice resulted in the production of more antiviral cytokines and resistance to RNA virus infection ^51^. A recent report showed that OTUD1 removes the K63-linked polyubiquitin chain from YAP, which is conjugated by SCF^Skp2^, and thus regulates the Hippo pathway and tumourigenesis^24^. Furthermore, OTUD1 reportedly down-regulates the IFN antiviral pathway by removing the K6-linked ubiquitin chain from IRF3 ^21^, and quite recently OTUD1 was found to suppress IBD by removing the K63 ubiquitin chain from RIP1^27^. Collectively, these findings indicate that OTUD1 functions as a crucial regulator of cancer progression, antiviral host defense responses, and inflammatory responses.

In this study, we initially characterized the N-terminal APGR as a disordered low complexity region (Fig. 1). APGR is required to suppress canonical NF-κB activation, exhibits a broad elution profile in gel filtration, and possesses an interaction site for KEAP1 (Figs. 1, 5). Therefore, APGR seems to play a crucial role in the physiological function of OTUD1.

After confirming that OTUD1 preferentially hydrolyzes K63-linked ubiquitin chains, we found that the K63-polyubiquitylations of NF-κB signalling factors, such as IRAK1, LUBAC subunits (HOIP and SHARPIN), NEMO, and RIP1, were increased upon inflammatory cytokine-stimulation in *OTUD1*-knockout cells, suggesting that these proteins are potential substrates of OTUD1 upon stimulation (Figs. 2, 7c). Since the ablation of *OTUD1* did not affect B cell receptor- and noncanonical NF-κB activation (Extended Data Fig. 3), OTUD1 principally regulates the inflammatory cytokine-induced canonical NF-κB pathway by hydrolysing K63 ubiquitin chains.

Although the TNF-α-induced NF-κB activation was upregulated in *OTUD1^-/-^*-cells, the activation of TBK1 and IKKε was suppressed (Extended Data Fig. 2), and in contrast to the previous report^21^, we determined that IRF3-mediated IFN antiviral signalling was down-regulated by the deficiency of *Otud1* (Fig. 3). Since TBK1 activation and IRF3 phosphorylation were suppressed in poly(dA:dT)- and poly(I:C)-treated *Otud1^-/-^*-MEFs, OTUD1 seems to be primarily involved in the upstream TBK1/IKKε activation (Fig. 7c), but not the downstream atypical K6-ubiquitylation of IRF3^21^, in the type I IFN production pathway.

The ubiquitylation and deubiquitylation of RIP1/RIP3 regulate the NF-κB activation, apoptotic, and necroptotic pathways^52^, and DUBs such as CYLD^53^, A20^54^, and OTUB1^55^ are involved in the regulation of the K63 ubiquitylation of RIP1/RIP3. In this study, we showed that the genetic deficiency of *OTUD1* accelerated the TNF-α-induced apoptotic and necroptotic cell death pathways, and the basal K63 ubiquitylation of RIP3 was enhanced in *Otud1^-/-^*-MEFs (Figs. 4, 7c), suggesting that RIP3 is an endogenous substrate of OTUD1 under physiological conditions. Although PELI1 and TRIM25 reportedly function as E3s to K48-linked ubiquitylation at Lys363 and Lys501 in RIP3, respectively, to induce proteasomal degradation^56, 57^, the E3s that catalyse the K63-ubiquitylation of RIP3 have not identified, although they may be involved in the regulation of human health and disease.

Importantly, we identified OTUD1 as a physiological interactor with KEAP1 through the ETGE motif in APGR (Figs. 5, 7c). KEAP1 senses ROS *via* critical Cys residues, and protects against oxidative and electrophilic stress as a component of the KEAP1-CUL3-RBX1 E3 complex to induce the proteasomal degradation of NRF2^39^. The DLG and ETGE motifs in the Neh2 domain of NRF2 are crucial for KEAP1-binding^41^, and the C-terminal Kelch domain of KEAP1 binds NRF2 as well as OTUD1. We showed that the K63-ubiquitylation of basal KEAP1 was upregulated in *Otud1^-/-^*-MEFs, indicating that Otud1 is physiological regulator of KEAP1. At present, several E3s, such as TRAF6^58^, TRIM25^59^, CKIP1, and SMURF1^60^, reportedly ubiquitylate NRF2-KEAP1. Therefore, OTUD1 may antagonise these E3s. Furthermore, since KEAP1 also binds with the phosphorylated ubiquitin-binding autophagy receptor protein p62/SQSTM1 through the KIR motif and regulates autophagy^61^, OTUD1 may regulate p62-KEAP1-mediated autophagy and related diseases, such as cancer progression.

In the presence of excess amounts of ROS, KEAP1 induces oxeiptosis, a caspase- and RIP3-independent apoptosis-like cell death, through binding with PGAM5 and dephosphorylating the evolutionally conserved Ser116 in AIFM1^40^. Moreover, upon TNF-α-induced necroptosis, PGAM5 binds necrosome, composed of RIP1-RIP3-MLKL, which mediate the activation of the mitochondrial fission factor Drp1 and mitochondrial fragmentation^62^. A recent report found that OTUD1 is involved in caspase-dependent and -independent apoptosis by facilitating the intranuclear translocation of AIFM1 and the degradation of MCL1^20^. The deubiquitylation of K244 in AIFIM1 by OTUD1 affects the mitochondrial structure and oxidative phosphorylation. We have shown that OTUD1 is a regulator of the KEAP1-PGAM5-AIFM1 axis, which affects ROS generation and cell death pathways upon oxidative and inflammatory responses (Figs. 5, 7c, Extended Data Fig. 5).

Free radicals, such as superoxides, hydroxyl radicals, peroxynitrite, and so on, induce oxidative damage, which causes genotoxicity, oxidations of proteins, lipids, and nucleotides, and cell death. Together, KEAP1-NRF2 and NF-κB co-operate the oxidative stress and inflammatory responses, and dysfunctions of these systems are associated with various disorders, such as neurodegeneration and inflammatory diseases^63, 64^. This study has demonstrated that OTUD1 is a central regulatory factor in these signal pathways, and the genetic ablation of *OTUD1* is associated with inflammatory bowel disease, hepatitis, sepsis, and kidney cancer (Figs. 6, 7, Extended Data Fig. 6)^21, 27^. Therefore, OTUD1 is a critical drug discovery target for the treatment of these diseases.

## Methods

### Reagents and plasmids

The following reagents were obtained as indicated: human TNF-α and IL-1β (BioLegend), poly(dA:dT) and high molecular weight poly(I:C) (Invivogen), LPS from *E. coli* O111:B4 (Invivogen), cycloheximide (Sigma-Aldrich), recombinant mouse CD40 ligand (R&D Systems), *α*-IgM (Southern Biotech), zVAD-FMK (ZVAD, Peptide Institute, Inc.), and necrostatin-1 (ENZO Life Sciences). The open reading frames of cDNAs, including 88 human DUBs, were amplified by reverse transcription-PCR. Mutants of these cDNAs were prepared by the QuikChange method, and the entire nucleotide sequences were verified. The cDNAs were ligated to the appropriate epitope sequences and cloned into the pcDNA3.1 (Invitrogen) and pGEX-6P-1 (GE Healthcare) vectors. For lentiviral transduction, pCSII-CMV-RfA-IRES-Blast (RIKEN BioResource Research Center) was used.

### Antibodies

The following antibodies were used for immunoblotting. Catalogue numbers and dilutions are indicated in parentheses. P-IκBα (#9246; 1:1,000), IκBα (#4812; 1:1,000), P-p105 (#4806; 1:1,000), p105 (#13586; 1:1,000), P-p65 (#3033; 1:1,000), p65 (#8242; 1:1,000), p100/p52 (#4882; 1:1,000), P-RelB (#4999; 1:1,000), RelB (#4922; 1:1,000), P-JNK (#4668; 1:1,000), JNK (#5668; 1:1,000), P-IRF3 (#4947; 1:1,000), IRF3 (#4302; 1:1,000), P-TBK1 (#5483; 1:1,000), TBK1 (#3504; 1:1,000), P-IKKε (#8766; 1:1,000), IKKε (#2690; 1:1,000), IκBζ (#9244; 1:1,000), caspase 3 (#9662; 1:1,000), cleaved caspase 3 (#9661; 1:1,000), caspase 8 (#9746; 1:1,000), caspase 8 (#4790; 1:1,000) cleaved caspase 8 (#9496; 1:1,000), PARP (#9542; 1:1,000), P-RIP1 (#31122; 1:1,000), P-RIP3 (#57220; 1:1,000), RIP3 (#15828; 1:1,000), P-MLKL (#37333; 1:1,000), MLKL (#37705; 1:1,000), IRAK1 (#4504; 1:1,000), CYLD (#8462; 1:1,000), OTULIN (#14127; 1:1,000), KEAP1 (#8047; 1:1,000), AIFM1 (#5318, 1:1,000), Tom20 (#42406, 1:1,000), Lamin A/C (#4777, 1:1,000), and GST (#2622, 1:1,000) were obtained from Cell Signaling. HOIL-1L (sc-393754; 1:250), TNFR1 (sc-7895; 1:1,000), VCAM (sc-8304; 1:1,000), and β-actin (sc-47778; 1:250) were purchased from Santa Cruz Biotechnology. HOIP (ab125189; 1:1,000), NEMO (ab178872; 1:3,000), and K63-linkage specific ubiquitin (ab179434; 1:1,000) were purchased from Abcam. RIP1 (610458; 1:1,000) and FADD (610399; 1:1,000) were from BD Transduction Laboratories. OTUD1 (NBP1-90484, Novus Biologicals; 1:1,000), tubulin (CLT9002, Cedarlane; 1:5,000), linear ubiquitin (Millipore, clone LUB9, MABS451; 1:1,000), SHARPIN (14626-1-AP, Proteintech; 1:5,000), KEAP1 (60027-1-Ig, Proteintech; 1:2,000), PGAM5(28445-1-AP, Proteintech; 1:1,000), DYKDDDDK (1E6, HRP-conjugate, Wako; 1:10,000), Myc (M192-7, HRP-conjugate, MBL; 1:20,000), and HA (11867423001, Roche; 1:2,000) were also used. To stimulate the lymphotoxin β receptor (LTβR), a 10:1 mix of the 4H8 WH2 (Adipogen, AG-20B-0008) and 3C8 (eBiosciences, 16-5671-82) monoclonal antibodies was used. For immunoprecipitation, Myc (sc-40, Santa Cruz Biotechnology; 1 μg), HA (16B12, BioLegend; 1 μg) and OTUD1 (NBP1-90484, Novus Biologicals; 3 μg) were used.

### Cell culture and transfection

HEK293T, HeLa (ATCC), and MEF cells were cultured in DMEM, containing 10% fetal bovine serum, 100 IU ml^-1^ penicillin G, and 100 μg ml^-1^ streptomycin, at 37°C under a 5% CO_2_ atmosphere. Bone marrow-derived macrophages (BMDM) were prepared basically as described^65^. Briefly, bone marrow cells were isolated from the femurs and tibias of B6 mice. After reticulocyte removal, the residual cells were cultured in 10% FCS/RPMI, supplemented with 5% of L-929 conditioned medium. The floating cells were removed, and the attached cells were passed every 2 days. The resultant adherent cells were used as macrophages on day 7. To obtain splenic B cells, suspensions of mouse splenocytes were prepared from freshly removed mouse spleens, and erythrocytes were depleted with ACK lysing buffer (0.15 M NH_4_Cl, 10 mM KHCO_3_, 0.1 mM Na_2_EDTA, pH 7.2), separated by BD IMag anti-mouse CD45R/B220 particles (BD Biosciences), and then maintained in RPMI containing 10% FBS, 55 μM 2-mercaptoethanol, and antibiotics, at 37 °C under a 5% CO_2_ atmosphere. Plasmid transfection into HEK293T cells was performed using Lipofectamine 2000 (Thermo Fisher Scientific) or polyethylenimine (PEI). To stimulate the cells, poly(dA:dT) or poly(I:C) was transfected using Lipofectamine 2000 (Thermo Fisher Scientific). For the stable expression of FLAG-tagged Otud1 in *Otud1*^-/-^-MEFs, lentiviral infection followed by selection with 5 µg ml^-1^ blasticidin was performed.

### Luciferase assay

HEK293T cells were cultured in 6-well or 24-well plates, and co-transfected with the pGL4.32 [*luc2P*/NF-κB-RE/Hygro] vector (Promega) and the phRL-TK *Renilla* Luciferase control reporter vector (Promega). At 24 h after transfection, the cells were lysed and the luciferase activity was measured with a GloMax 20/20 luminometer (Promega), using the Dual-Luciferase Reporter Assay System (Promega).

### Screening for DUBs

HEK293T cells cultured in 6-well plates were transfected with pcDNA3.1 plasmids encoding DUB (1 μg), HA-HOIP (30 ng), HOIL-1L-HA (15 ng), and HA-SHARPIN (10 ng), with NF-κB and control luciferase reporters. After 24 h, the luciferase activity was measured as described above. Taking the NF-κB-luciferase activity induced by LUBAC alone as 100%, the relative residual activity in the presence of the respective DUBs was calculated.

### Gel filtration analysis

Cells were suspended in 50 mM Tris-HCl, pH 7.5, containing 1 mM MgCl_2_, 1 mM DTT, 1 mM PMSF, and a protease inhibitor cocktail (Complete EDTA-free, Roche), and homogenized by a Dounce homogenizer. After adding an equal volume of buffer containing 300 mM NaCl, lysates were centrifuged at 100,000 *g* for 30 min. The supernatant was fractionated on an ÄKTA pure 25 chromatography system using a Superdex 200 HR (10/30) column (Cytiva), which was equilibrated with 50 mM Tris-HCl, pH 7.5, containing 150 mM NaCl. The column was calibrated by the elution profile of a Gel Filtration HMW Calibration Kit (Cytiva).

### Preparation of recombinant OTUD1

The pGEX-6P-1 expression vector, which encodes glutathione *S*-transferase (GST)-fused OTUD1-Wt, was expressed in *E. coli* BL21(DE3) (Novagen), and the recombinant protein was prepared as described previously^66^.

### *In vitro* DUB assay

For the *in vitro* DUB assay with diubiquitins, HEK293T cells were transfected with FLAG-OTUD1-Wt or catalytically inactive FLAG-OTUD1-CA and lysed in lysis buffer, containing 20 mM Tris-HCl, pH 7.5, 150 mM NaCl, 1% Triton X-100, and 1 mM DTT. The cell lysates were immunoprecipitated by a rabbit anti-FLAG antibody (Sigma-Aldrich, F7425) and protein A beads (Cytiva). After a wash with lysis buffer, the beads were reacted with 0.5 μg of M1-, K6-, K11-, K27-, K29-, K33-, K48-or K63-linked diubiquitins (Boston Biochem) in 20 mM Tris-HCl, pH 7.5, containing 1 mM DTT, for 30 min at 37 °C. For the *in vitro* deubiquitylation assay with polyubiquitin chains, the M1-, K48-, or K63-linked polyubiquitin chains (2.0 μg), prepared as described previously^14^, were treated with 0.6 μM recombinant GST-OTUD1-Wt in a total volume of 20 μl of 20 mM Tris-HCl, pH 7.4, containing 1 mM DTT, for the indicated time periods at 37 °C.

### Immunoprecipitation, SDS-PAGE, and immunoblotting

Cells were lysed with 50 mM Tris-HCl, pH 7.5, containing 150 mM NaCl, 1% Triton X-100, 2 mM PMSF, and complete protease inhibitor cocktail (Sigma). Immunoprecipitation was performed using appropriate antibodies followed by Protein G agarose beads (Cytiva) at 4°C with gentle rotation. Immunoprecipitates were washed five times with the lysis solution. Samples were separated by SDS-PAGE and transferred to PVDF membranes. After blocking, the membranes were incubated with the appropriate primary antibodies, followed by an incubation with HRP-conjugated secondary antibodies. The chemiluminescent images were obtained with an LAS4000 imaging analyzer (GE Healthcare) or a Fusion Solo S imaging system (Vilber).

### Construction of *OTUD1*-knockout cells

To generate *OTUD1*-knockout HeLa and HEK293T cells, the gRNA cloning vector (Addgene; #41824) targeting 5ʹ-CCGagaccggtgagtgccagccc-3ʹ and 5ʹ-tacccggagttgctggccaTGG-3ʹ (Capital letters indicate PAM-sequence) in the coding sequence of human *OTUD1*, and the pCAG-hCas9 expression vector (Addgene; #51142) were transfected using Lipofectamine 3000 (Thermo Fisher). After 96 h, cells were cloned by limiting dilution to obtain single cell clones. The clones were validated by *Age*I or *Nco*I digestion, sequenced with PCR-amplified fragments to confirm mutations, and subjected to immunoblotting with an anti-OTUD1 antibody.

### Construction of *Otud1*-knockout mice

C57BL/6J mice were used as the source of embryos for micromanipulation and subsequent breeding trials. A mixture of sgRNAs targeting the 5ʹ-UTR of *Otud1*: 5ʹ-attcgcggctcctgacgcggCGG-3ʹ (Capital letters indicate PAM-sequence) and 3ʹ-UTR of *Otud1*: 5ʹ-gaaccacacgaactctaattAGG-3ʹ (Capital letters indicate PAM-sequence), and Cas9 protein was injected into the cytoplasm of one cell-stage fertilized embryos. Zygotes that survived were transferred into the oviducts of pseudopregnant foster mothers. The heterozygous mutant mice were intercrossed to produce the homozygous mutant mice. For PCR primers to detect the wild-type *Otud1* allele, we used 5ʹ-GTATTTGGCTCAGTTGGCTC-3ʹ and 5ʹ-TTCATACCGTGAGACATCAG-3ʹ oligo DNAs to detect the 450 bp band. For the *Otud1* deleted allele, we used 5ʹ-ATCCGCGCGCTGCACTCCTG-3ʹ and 5ʹ-TTCATACCGTGAGACATCAG-3ʹ oligo DNAs to detect the 440 bp band (which was 1,960 bp in the wild-type allele). C57BL/6J mice were kept in a specific pathogen-free, temperature-controlled facility at the Graduate School of Medicine, Osaka City University. Mice were euthanized according to the guidelines of the Animal Experiment Committee of Osaka City University, and all efforts were made to minimize the suffering of the animals. MEFs were prepared from littermate embryos (E12.5) of Wt- and *Otud1*-KO mice, transfected with an SV40 T-antigen containing pEF321-T plasmid, and then immortalized.

### K63- and M1-TUBE pulldowns

Cells were lysed in RIPA buffer (50 mM Tris-HCl, pH 7.5, containing 150 mM NaCl, 1% NP-40, 1% sodium deoxycholate, and 0.1% SDS) supplemented with 2 mM PMSF, protease inhibitor cocktail, and 10 mM *N*-ethylmaleimide, and centrifuged at 20,000 *g* for 20 min at 4 °C. The cleared lysates (4 mg) were diluted 10-fold with 1% Triton X-100 buffer (50 mM Tris-HCl, pH 7.5, 150 mM NaCl, 1% Triton X-100), and then incubated with 30 μl of K63-Tandem Ubiquitin Binding Entity (TUBE)-Agarose (LifeSensors, UM401), or 2.4 μg of K63-TUBE-Biotin (LifeSensors, UM304) or M1-TUBE-Biotin (LifeSensors, UM306) with 30 μl of Dynabeads M-280 Streptavidin (Veritas), overnight at 4°C. The precipitates were washed five times with 1% Triton X-100 buffer, boiled in SDS sample buffer, and then subjected to immunoblotting analysis.

### qPCR

Cell lysis, reverse-transcription, and qPCR were performed with a SuperPrep Cell Lysis RT Kit for qPCR (TOYOBO) and Power SYBR Green PCR Master Mix (Life Technologies), according to the manufacturers’ instructions. Quantitative real-time PCR was performed with a Step-One-Plus PCR system (Applied Biosystems) by the ΔΔCT method, using the following oligonucleotides: human *IL-6* sense, 5’-AGCCACTCACCTCTTC-3’, and human *IL-6* anti-sense, 5’-GCCTCTTTGCTGCTTT-3’; human *ICAM1* sense, 5’-GTGGTAGCAGCCGCAGT-3’ and human *ICAM1* anti-sense, 5’-TTCGGTTTCATGGGGGT-3’; human *BIRC3* sense, 5’-AGATGAAAATGCAGAGTCATCAAT-3’ and human *BIRC3* anti-sense, 5’-CATGATTGCATCTTCTGAATGG-3’; and human *GAPDH* sense, 5’-AGCAACAGGGTGGTGGAC-3’ and human *GAPDH* anti-sense, 5’-GTGTGGTGGGGGACTGAG-3’.

For analyses with mouse-derived immune cells or tissues, total mRNA was extracted with an RNeasy Mini Kit (QIAGEN), and then transcribed to cDNA with ReverTra Ace qPCR RT Master Mix with gDNA Remover (TOYOBO), according to the manufacturers’ instructions. Quantitative real-time PCR was performed with Power SYBR Green PCR Master Mix (Life Technologies) and a Step-One-Plus PCR system by the ΔΔCT method, using the following oligonucleotides: mouse *Ifnb1* sense: 5’-CCCTATGGAGATGACGGAGA-3’ and mouse *Ifnb1* anti-sense: 5’-CTGTCTGCTGGTGGAGTTCA-3’; mouse *Isg15* sense: 5’-GGAACGAAAGCGGCCACAGCA-3’ and mouse *Isg15* anti-sense: 5’-CCTCCATGGGCCTTCCCTCGA-3’; mouse *Isg56* sense: 5’-CCCTATGGAGATGACGGAGA-3’ and mouse *Isg56* anti-sense: 5’-CTGTCTGCTGGTGGAGTTCA-3’; mouse *Cxcl10* sense, 5’-CCAAGTGCTGCCGTCATTTTC-3’ and mouse *Cxcl10* anti-sense, 5’-GGCTCGCAGGGATGATTCAA-3’; mouse *Tnfaip3* sense, 5’-GCTTTCGCAGAGGCAGTAACAG-3’ and mouse *Tnfaip3* anti-sense, 5’-AGCAAGTGCAGGAAAGCTGGCT-3’; mouse *Bcl-xl* sense, 5’-GGTGAGTCGGATTGCAAGTT-3’ and mouse *Bcl-xl* anti-sense, 5’-GCTGCATTGTTCCCGTAGAG-3’; mouse *Hmox-1* sense, 5’-AAGCCGAGAATGCTGAGTTCA-3’ and mouse *Hmox-1* anti-sense, 5’-GCCGTGTAGATATGGTACAAGGA-3’; mouse *Nfe2l2* sense, 5’-TCTTGGAGTAAGTCGAGAAGTGT-3’ and mouse *Nfe2l2* anti-sense, 5’-GTTGAAACTGAGCGAAAAAGGC-3’; mouse *Tnf* sense, 5’-TAGCCAGGAGGGAGAACAGA-3’ and mouse *Tnf* anti-sense, 5’-TTTTCTGGAGGGAGATGTGG-3’; mouse *Il-1b* sense, 5’-CCCTGCAGCTGGAGAGTGTGGA-3’ and mouse *Il-1b* anti-sense, 5’-TGTGCTCTGCTTGTGAGGTGCTG-3’; and mouse *Gapdh* sense, 5’-TTTGGCATTGTGGAAGGGCTCAT-3’ and mouse *Gapdh* anti-sense, 5’-CACCAGTGGATGCAGGGATGATGT-3’.

### Analyses of TNFR complexes I and II

The TNFR complex I analysis was performed as described previously^14^. Briefly, parental and *OTUD1^-/-^*-HeLa cells were stimulated for the indicated times with 1 μg ml^-1^ FLAG-tagged TNF-α, and then lysed in 1 ml lysis buffer, containing 50 mM Tris-HCl, pH 7.5, 150 mM NaCl, 1% Triton X-100, and complete protease inhibitor cocktail. After centrifugation at 20,000 *g* for 15 min, the supernatants were immunoprecipitated with 40 μl anti-FLAG M2 beads (Sigma) overnight at 4°C. For the TNFR complex II analysis, parental and *OTUD1^-/-^*-293T cells were treated with 10 ng ml^-1^ TNF-α and 10 μg ml^-1^ CHX. The cell lysates were then immunoprecipitated with an anti-FADD antibody (Santa Cruz; sc-271748) and immunoblotted.

### RNA-seq

Parental and *OTUD1^-/-^*-HeLa cells stimulated with 1 ng ml^-1^ IL-1β for 1 and 3 h, and *Otud1^+/+^*- and *Otud1^-/-^*-MEFs stimulated with 10 μg ml^-1^ poly(I:C) for 2 and 4 h, were lysed, and the total RNA was extracted using a RNeasy Mini kit (Qiagen, 74104) according to the manufacturer’s instructions. The mRNA template library was constructed with a TruSeq Stranded mRNA LT Sample Prep Kit (Illumina), and the sequencing analysis was performed by Macrogen Corp. (Korea), using a NovaSeq 6000 sequencer (Illumina) and an S4 Reagent Kit. The data were processed by a Multidimensional Scaling Analysis and a Hierarchical Clustering Analysis by the Euclidean algorithm with the Complete Linkage program.

### Cell survival assay

The number of viable cells was measured with a CellTiter-Glo Luminescent Cell Viability Assay (Promega), which quantifies the ATP content, and a trypan blue exclusion assay. The cell viability was also continuously monitored as an impedance-based cell index, using an xCELLigence RTCA S16 instrument (ACEA Biosciences Inc.). For the experiments, the parental and *OTUD1*^-/-^-HEK293T cells (40,000 cells/well) or MEF cells (20,000 cells/well) were seeded in an E-Plate VIEW 16 (ACEA Biosciences, Inc.). The next day, the cells were stimulated with TC or TCZ, and the cell index was continuously monitored every 15 min. The data were analysed with real-time cell analysis (RTCA) software, and normalized against the cell indices at the time of TC or TCZ treatment.

### ROS measurement

Intracellular ROS levels were measured using an ROS Assay Kit-Highly Sensitive DCFH-DA (Dojindo), which is converted to the highly fluorescent 2′,7′-dichlorofluorescein (DCF) in the presence of an oxidant, according to the manufacturer’s instructions. Briefly, cells were cultured in 96-well black plates (clear bottom) and treated as indicated. After treatment, the cells were washed with HBSS (Gibco), and incubated with the DCFH-DA solution for 30 min at 37°C under a 5% CO_2_ atmosphere. ROS production was analysed with a Varioskan LUX multimode microplate reader (Thermo Fisher Scientific) with excitation and emission wavelengths at 490 and 520 nm, respectively.

### Identification of OTUD1-associated proteins by LC-MS/MS

An empty vector or a FLAG-tagged OTUD1 expressing vector was transfected into HEK293T cells in 10-cm dishes. At 24 h after transfection, the cells were washed with HEPES-saline (20 mM HEPES-NaOH, pH 7.5, 137 mM NaCl) and lysed with 20 mM HEPES-NaOH, pH 7.5, 150 mM NaCl, 1% Triton X-100, and complete protease inhibitor cocktail (Sigma). After centrifugation at 20,000 *g*, 4°C for 10 min, the supernatants were incubated with anti-FLAG M2 Magnetic Beads (M8823, Sigma) for 1 h at 4°C with gentle rotation. The beads were collected using a magnetic stand and washed four times with the 1% Triton X-100 lysis buffer, and then another three times with 50 mM ammonium bicarbonate. Proteins on the beads were digested with 200 ng trypsin/Lys-C mix (Promega, V5072) for 16 h at 37°C. The digests were reduced, alkylated, acidified with trifluoroacetic acid (Wako, 206–10731; TFA), and desalted using GL-Tip SDB (GL Sciences, 7820–11200). The eluates were evaporated in a SpeedVac concentrator and then dissolved in 0.1% TFA and 3% acetonitrile (ACN). The LC-MS/MS analysis of the resultant peptides was performed on an EASY-nLC 1200 UHPLC connected to an Orbitrap Fusion mass spectrometer through a nanoelectrospray ion source (Thermo Fisher Scientific). The peptides were separated on a 75 μm inner diameter × 150 mm C18 reversed-phase column (Nikkyo Technos) with a linear 4–32% ACN gradient for 0–100 min, followed by an increase to 80% ACN for 10 min and a final hold at 80% ACN for 10 min. The mass spectrometer was operated in the data-dependent acquisition mode with a maximum duty cycle of 3 s. MS1 spectra were measured with a resolution of 120,000, an automatic gain control (AGC) target of 4e5, and a mass range from 375 to 1,500 *m/z*. HCD MS/MS spectra were acquired in the linear ion trap with an AGC target of 1e4, an isolation window of 1.6 *m/z*, a maximum injection time of 35 ms and a normalized collision energy of 30. Dynamic exclusion was set to 20 s. Raw data were directly analysed against the SwissProt database restricted to *Homo sapiens*, using Proteome Discoverer version 2.4 (Thermo Fisher Scientific) with the Mascot search engine. The search parameters were as follows: (a) trypsin as an enzyme with up to one missed cleavage; (b) precursor mass tolerance of 10 ppm; (c) fragment mass tolerance of 0.6 Da; (d) carbamidomethylation of cysteine as a fixed modification; and (e) acetylation of protein N-terminus and oxidation of methionine as variable modifications. Peptides and proteins were filtered at a false discovery rate (FDR) of 1%, using the Percolator node and the Protein FDR Validator node, respectively. Label-free quantification was performed based on the intensities of precursor ions, using the Precursor Ions Quantifier node. Normalization was performed such that the total sum of abundance values for each sample over all peptides was the same.

### PRM analysis

To quantify KEAP1, AIFM1, CUL3, and PGAM5, two or three peptides/protein were measured by parallel reaction monitoring (PRM), an MS/MS-based targeted quantification method using high-resolution MS. The LC-MS/MS analysis was performed on an EASY-nLC 1200 UHPLC connected to a Q Exactive Plus mass spectrometer through a nanoelectrospray ion source (Thermo Fisher Scientific). Targeted HCD MS/MS scans were acquired by a time-scheduled inclusion list at a resolution of 70,000, an AGC target of 2e5, an isolation window of 2.0 *m/z*, a maximum injection time of 300 ms, and a normalized collision energy of 27. Time alignment and relative quantification of the transitions were performed using the Skyline software.

### Sendai virus infection

MEFs and BMDMs were infected with Sendai virus (SeV) strain Cantell at a cell infectious unit (CIU) of 10 for 1 h. The washed cells were then incubated for the indicated times.

### Dextran sulfate sodium (DSS)-induced IBD model

To analyse the DSS-induced colitis model, 8-week old Wt- and *Otud1^-/-^*-C57/BL/6 background male mice were used. Each mouse was weighed once a day and monitored for the appearance of bloody and loose stools. Mice were sacrificed after a 7 day administration of 2.5% DSS (MW 36,000-50,000, MP Biomedicals) in drinking water. The stool consistency (diarrhea) and visible fecal blood were scored separately on a scale of 0 to 3, as described^67^

### Acute hepatitis model mice

LPS with D-galactosamine (GalN)-induced acute hepatitis model mice were prepared by intraperitoneal injections of LPS (10 μg kg^-1^) and GalN (800 mg kg^-1^, Nacalai Tesque) in 200 μl PBS into 8-week old Wt- and *Otud1^-/-^*-C57/BL/6 background mice. For the survival test, mice were injected with LPS/GalN and monitored every 15 min for 10 h. For immunoblotting or histopathologic examination, mice were sacrificed at 5 h after injection, and liver tissues were lysed with RIPA buffer or fixed in a 4% paraformaldehyde (PFA) solution.

### Sepsis model mice

LPS (20 mg kg^-1^) was intraperitoneally injected into 8-week old Wt- and *Otud1^-/-^*-C57/BL/6 background mice, and their survival was monitored.

### Histopathological analysis

In the DSS-induced colitis model, colons and spleens were excised. Colons were distended in 0.9% saline and opened longitudinally, and then cut into three segments (proximal, middle, and distal colon). A transverse section of the spleen was made at the largest extension of the organ. After fixation in a 4% PFA solution, colon and spleen specimens were embedded in paraffin and processed for hematoxylin/eosin (H&E) and immunohistochemical staining. The histologic scores of colitis were evaluated using a quantitative evaluation form, as described^68^.

In the sepsis model or acute hepatitis model, a total of 5 sections of liver tissue (one section each from the left lateral lobe, left middle lobe, right middle lobe, right lateral lobe and caudate lobe) were fixed in a 4% PFA solution, embedded in paraffin, and processed for H&E and immunohistochemical staining.

### Immunohistochemical analysis

Paraffin sections (4-µm thickness) were deparaffinized and dehydrated through a graded ethanol series. 8-Hydroxy-2-deoxy-guanosine (8-OHdG) staining was performed using a Histofine MOUSESTAIN kit (Nichirei, 414321) and an anti-8-OHdG (8-hydroxy-2’-deoxyguanosine) antibody (JaICA, MOG-100P) diluted 1:50. To detect cell death (necrosis and/or apoptosis), TUNEL staining was performed using an ApopTag® Peroxidase *In Situ* Apoptosis Detection Kit (Millipore, S7100), according to the manufacturer’s instructions. TUNEL-positive cells in the spleens were counted in six 40x fields of each specimen, and quantified as the number of positive cells per field. Hepatocellular death was scored on a semiquantitative scale of 1 to 3, using the following criteria: 1, **≤**1% TUNEL-positive cells; 2, >1% but **≤**5% TUNEL-positive cells; 3, >5% TUNEL-positive cells. Microscopic image capture was performed using an Olympus BX53 microscope equipped with a DP72 digital microscope camera and a DP2-BSW image acquisition system (Olympus Corp., Tokyo, Japan).

### Gene expression database analysis

The correlation of patient survival and the level of OTUD1 gene expression, and the OTUD1 gene comparison in normal or tumour kidney cortex, were analysed online using GEPIA (http://gepia.cancer-pku.cn/). For the analysis of gene expression correlation, TCGA tumour (KIRC), TCGA normal (KIRC), and GTEx (Kidney-Cortex) were selected.

### Statistics

One-way ANOVA followed by a post-hoc Tukey HSD test, *t*-test, Mann-Whitney test, and Log-rank test of Kaplan-Meier survival curve were performed using the GraphPad Prism 8 software. For all tests, a *P* value of less than 0.05 was considered statistically significant.

## Acknowledgements

We thank Dr. Kohei Nishino (Tokushima Univ.) for MS analysis, Dr. Masaya Fukushi (Hiroshima Univ.) for the generous gift of SeV, Prof. Kohsuke Takeda (Nagasaki Univ.) for helpful discussions on oxeiptosis, Dr. Eiji Goto, Dr. Takanori Abe, Dr. Seigo Terawaki, Ms. Wakana Koeda, Ms. Shiori Motoyama, Ms. Chihiro Yamada, Ms. Yukimi Kira (Osaka City Univ.), and Dr. Yuki Katayama (Gunma Univ.) for technical assistance, and the Research Support Platform of Osaka City University Graduate School of Medicine for technical assistance. This work was partly supported by AMED under Grant Number JP21gm6410013 (D.O.), and a Grant for Research Program on Hepatitis from the Japan Agency for Medical Research and Development (19fk0210050h0001 to F.T.), MEXT/JSPS KAKENHI grants (Nos. JP16H06276 (AdAMS) and JP21H02688 to F.T., JP21K06873, JP21H00291, and JP 20H05337 to D.O., and JP20K16146 to K.S.), JST ACT-X Grant Number JPMJAX2117 (K.S.), and by the Takeda Science Foundation (F.T. and D.O.), Kobayashi Foundation (F.T.), and Joint Usage and Joint Research Programs of the Institute of Advanced Medical Sciences, Tokushima University (F.T.).

## Author contributions

D.O., K.S., H.T., and K.K. performed biochemical and cell biological experiments, M.G. performed histochemical analyses, H.K. conducted mass spectrometric analyses, and M.S. and S.H. participated in the mouse analyses. H.W., D.T., T.S., and F.T. conceived the project. D.O., M.G., H.K., K.S., and F.T. wrote the manuscript and all of the authors participated in commenting on the manuscript.

## Conflict of interest

The authors declare no competing interests.

## Notes

### Competing Interest Statement

The authors have declared no competing interest.

