## Supplementary Information for "OTUD1 deubiquitylase regulates NF-κB- and KEAP1-mediated inflammatory responses and reactive oxygen species-associated cell death pathways"

**Extended Data Table 1** Histopathological changes in the livers of mice 24 h after LPS treatment.

| | LPS | Inflammatory cell infiltration | Fatty change | Hydropic degeneration | Midzonal hepatocellular necrosis |
|---|---|---|---|---|---|
| Male | | | | | |
| *Otud1*$^{+/+}$ | - | 0/3 | 0/3 | 0/3 | 0/3 |
| | + | 6/6 | 6/6 | 2/6 | 0/6 |
| *Otud1*$^{-/-}$ | - | 0/3 | 0/3 | 0/3 | 0/3 |
| | + | 6/6 | 6/6 | 0/6 | 0/6 |
| Female | | | | | |
| *Otud1*$^{+/+}$ | - | 0/3 | 0/3 | 0/3 | 0/3 |
| | + | 6/6 | 0/6 | 4/6 | 2/6 |
| *Otud1*$^{-/-}$ | - | 0/3 | 0/3 | 0/3 | 0/3 |
| | + | 6/6 | 0/6 | 3/6 | 5/6 |

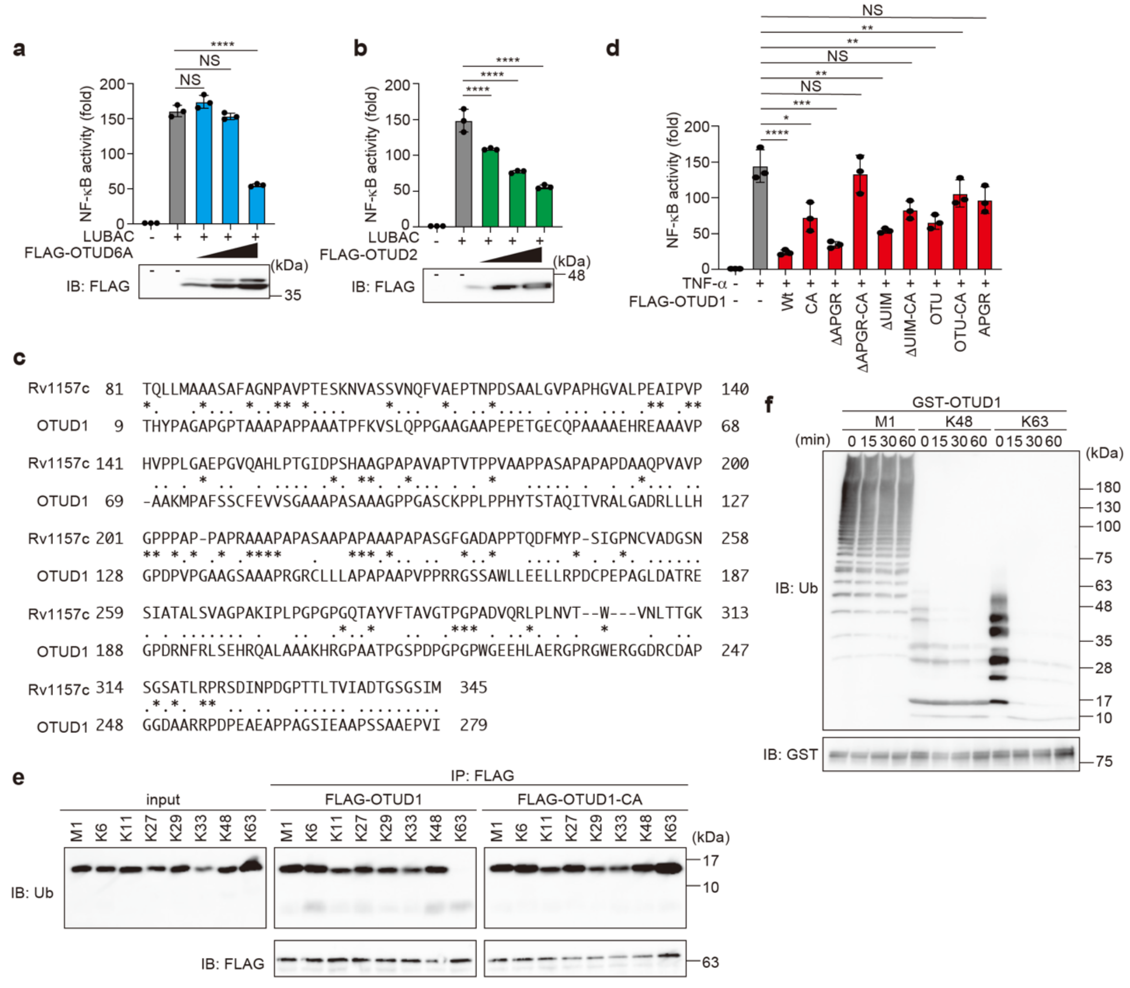


**Extended Data Fig. 1 Characterization of LUBAC-suppressive DUBs, including OTUD1. a, b,** OTUD6A and OTUD2 suppress LUBAC activity. The effects of increasing amounts (0.1, 0.3, and 1.0 µg) of OTUD6A (**a**) or OTUD2 (**b**) on LUBAC-induced NF-κB activity were analysed, as in Fig. 1b. **c,** Amino acid sequence alignment of hypothetical protein Rv1157c in *Mycobacterium tuberculosis* and the N-terminal region of OTUD1. Identical and chemically similar residues are denoted by *asterisks* and *dots*, respectively. **d,** Effect of Wt- and mutants of OTUD1 on the TNF-α-induced NF-κB activity. A similar analysis to that in Fig. 1d was performed by stimulating cells with 10 ng $ml^{-1}$ TNF-α for 6 h. **a, b, d,** Data are shown as mean ± SD by ANOVA post-hoc Tukey test (*n* = 3). *: *P*<0.05, **: *P*<0.01, ***: *P*<0.001, ****: *P*<0.0001, NS: not significant. **e,** OTUD1 hydrolyzes K63-diubiquitin. Wt or the active-site mutant of OTUD1 was expressed in HEK293T cells, and immunoprecipitated with anti-FLAG antibody-immobilized beads. Various diubiquitins were then treated with the immunoprecipitates for 1 h, and the cleavage of diubiquitin was investigated by immunoblotting with the indicated antibodies. **f**, OTUD1 cleaves K63-linked polyubiquitin chains. *In vitro* DUB assays were performed using recombinant GST-OTUD1 and M1-, K48-, and K63-linked polyubiquitin chains. Samples were incubated for the indicated periods and immunoblotted with the indicated antibodies.

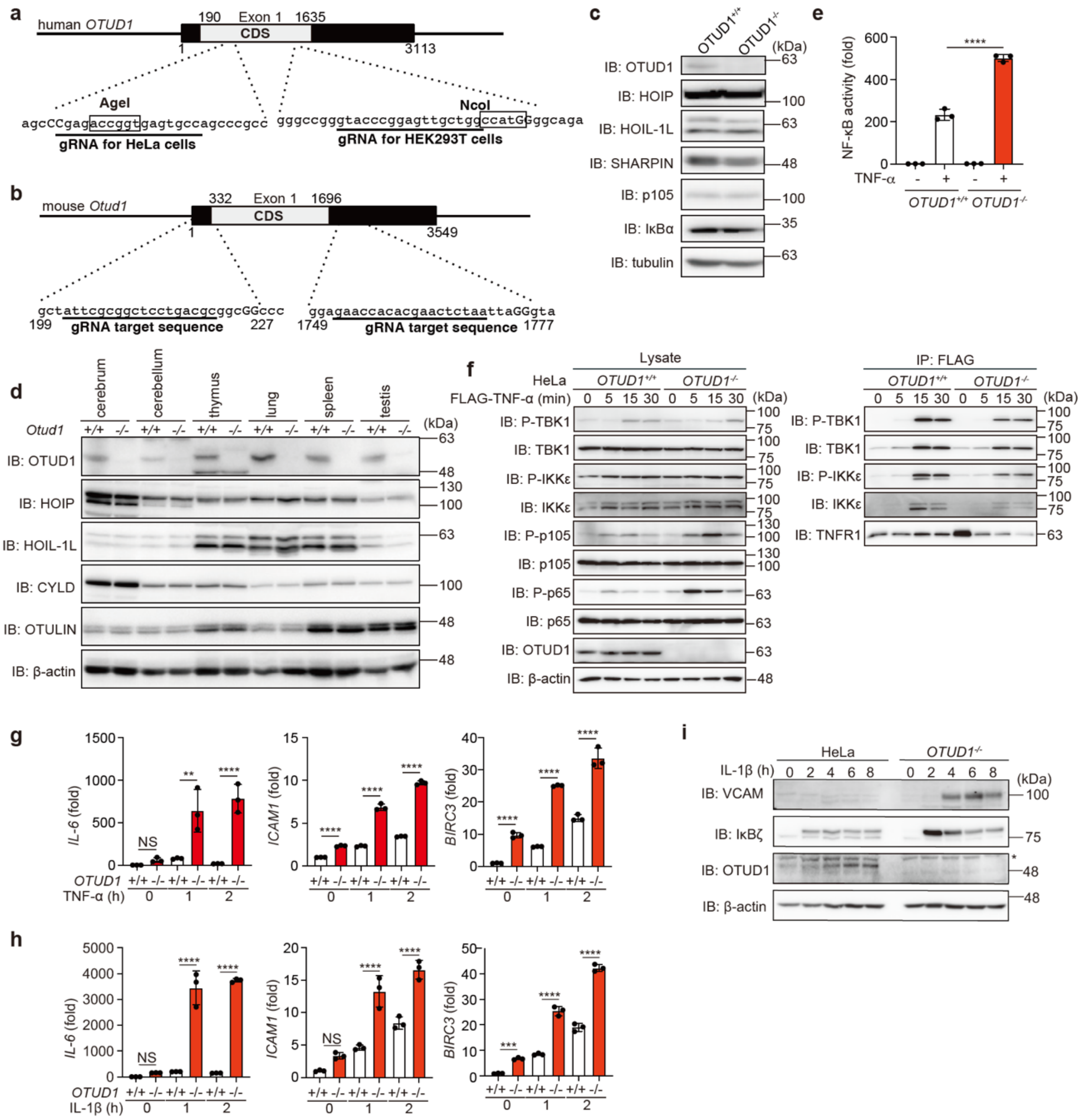


**Extended Data Fig. 2 Construction of *OTUD1*-deficient cells and mice, and their effects on inflammatory cytokine-induced NF-κB activation. a**, Scheme for *OTUD1*-KO targeting. Two independent gRNAs were used to target exon 1 of the *OTUD1* gene in HeLa and HEK293T cells. **b**, Scheme for *Otud1*-KO targeting in mouse. Two gRNAs were used to delete the entire exon 1 of the *Otud1* gene. **c**, Immunoblotting of parental and *OTUD1*$^{-/-}$-HEK293T cells. The cell lysates were immunoblotted with the indicated antibodies. **d**, Expression of Otud1 in various mouse tissues. Immunoblotting analyses of lysates from various *Otud1*$^{+/+}$- and *Otud1*$^{-/-}$-mouse tissues were performed. **e**, Enhanced TNF-α response in *OTUD1*-deficient cells. The NF-κB luciferase reporter was transfected into parental- and *OTUD1*$^{-/-}$-HEK293T cells, and luciferase activity was analysed after 6 h stimulation with 60 ng ml$^{-1}$ TNF-α. **f,** Reduced TNF-α-mediated recruitment and activation of TBK1/IKKε to TNFR. A similar analysis as in Fig. 2a was performed, and samples were immunoblotted with the indicated antibodies. **g, h,** Enhanced expression of NF-κB target genes in inflammatory cytokine-treated *OTUD1*$^{-/-}$-HeLa cells. Parental and *OTUD1*$^{-/-}$-HeLa cells were stimulated with 10 ng ml$^{-1}$ TNF-α (**g**) or 1 ng ml$^{-1}$ IL-1β (**h**) for the indicated periods, and a qPCR analysis was performed. **e**, **g**, **h**, Data are shown as mean ± SD by ANOVA post-hoc Tukey test ($n$ = 3). **: $P$<0.01, ***: $P$<0.001, ****: $P$<0.0001, NS: not significant. **i,** Increased expression of NF-κB-target proteins in *OTUD1*$^{-/-}$-HeLa cells. Cells were stimulated with 1 ng ml$^{-1}$ IL-1β for the indicated periods, and cell lysates were analysed by immunoblotting. *: nonspecific signal.

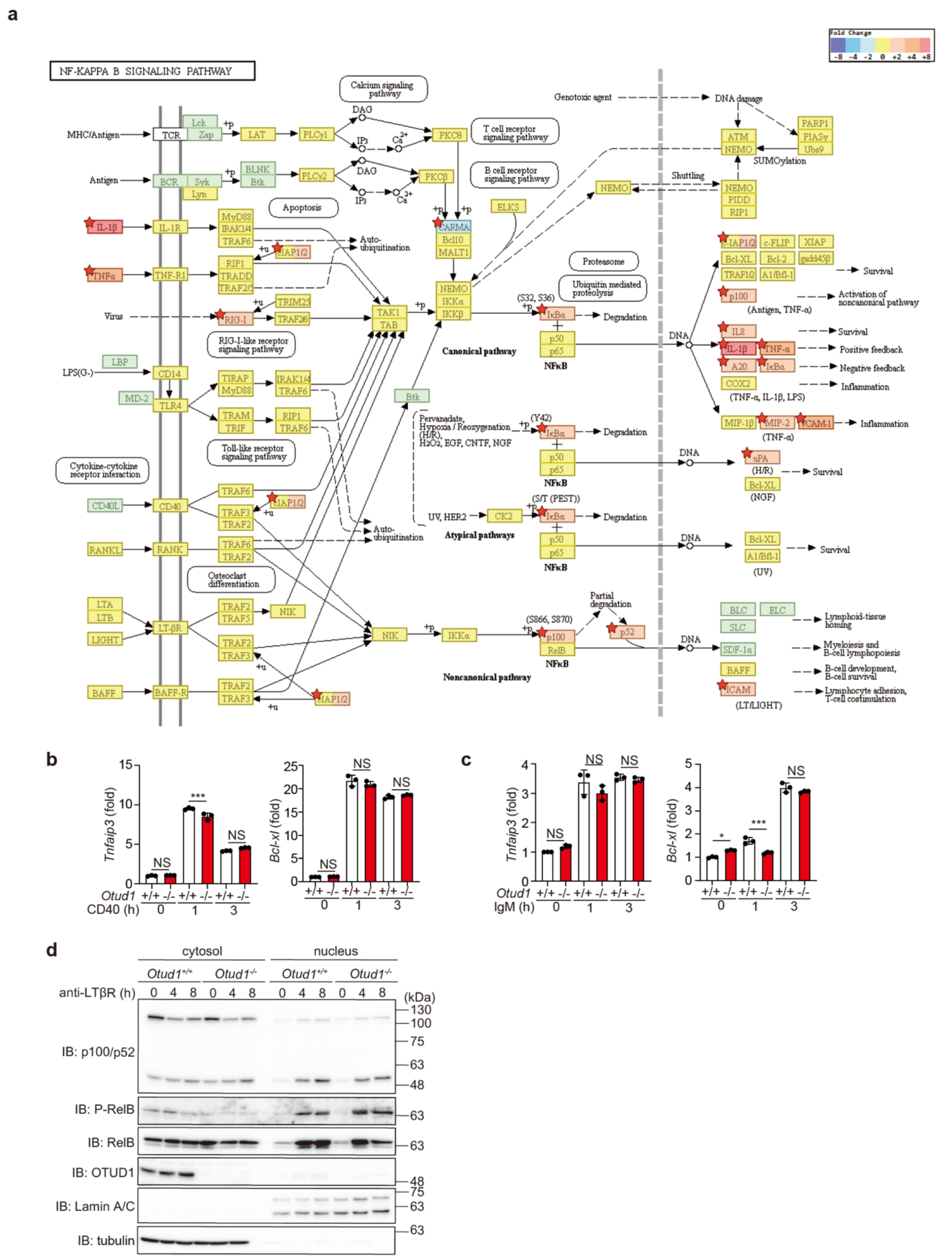

**Extended Data Fig. 3 Effect of OTUD1 on NF-κB signalling pathways. a**, Enhanced expression of NF-κB target genes in IL-1β-treated *OTUD1*$^{-/-}$ cells. The KEGG NF-κB signalling pathway map of various genes in IL-1β-treated parental and *OTUD1*$^{-/-}$-HeLa cells is shown. **b**, **c**, The genetic deficiency of

*Otud1* does not affect CD40- and B cell receptor-mediated NF-κB activation. Splenic B cells from *Otud1*$^{+/+}$- and *Otud1*$^{-/-}$-mice were stimulated with 3 μg $ml^{-1}$CD40L (**b**) or 5 μg $ml^{-1}$ IgM (**c**) for the indicated periods, and qPCR analyses were performed. Data are shown as mean ± SD ($n$ = 3). *: $P$<0.05, ***: $P$<0.001, NS: not significant. **d**, OTUD1 is not involved in the non-canonical NF-κB pathway. *Otud1*$^{+/+}$- and *Otud1*$^{-/-}$-MEFs were stimulated with 0.3 μg $ml^{-1}$ anti-lymphotoxin β receptor (LTβR) antibodies for the indicated periods. Cell lysates were separated into cytosolic and nuclear fractions, and immunoblotted with the indicated antibodies.

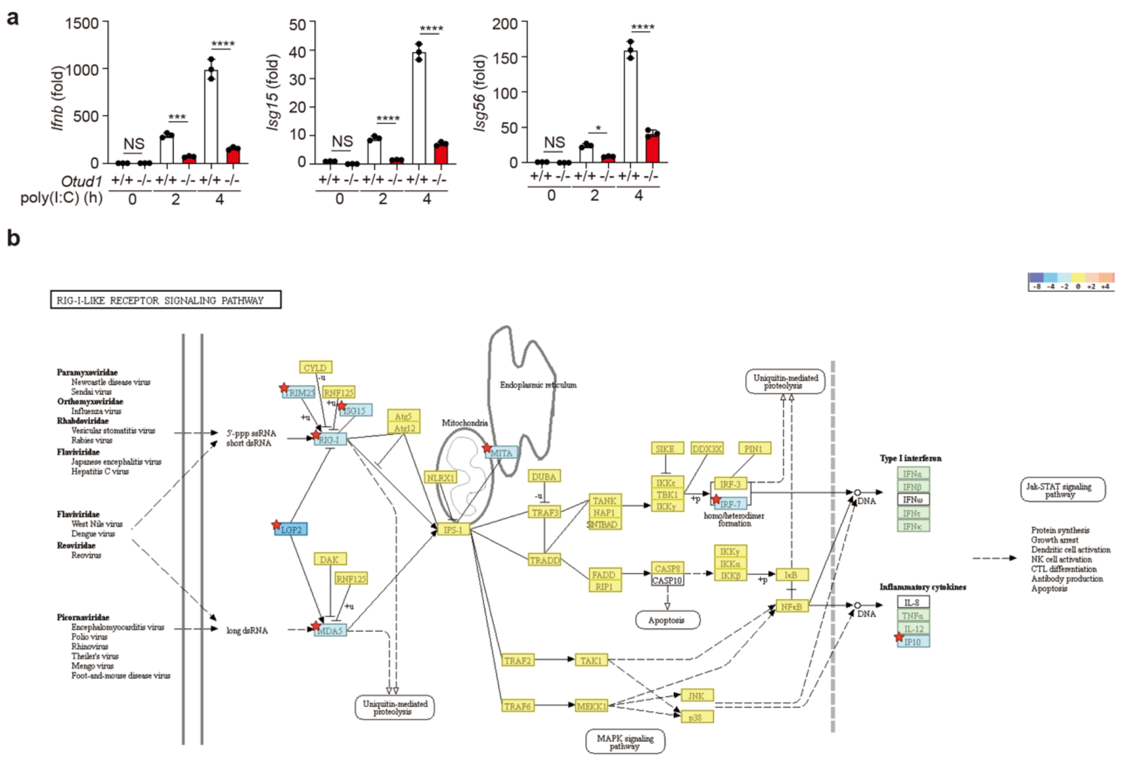


**Fig. 4 Reduced antiviral signalling in *Otud1*$^{-/-}$-MEFs. a**, *Otud1*$^{+/+}$- and *Otud1*$^{-/-}$-MEFs were stimulated with 10 μg ml$^{-1}$ poly(I:C) for the indicated periods, and qPCR was performed. Data are shown as mean ± SD ($n$ = 3). *: $P$<0.05, ***: $P$<0.001, ****: $P$<0.0001, NS: not significant. **b,** Reduced IFN antiviral signalling in *Otud1*$^{-/-}$-MEFs. KEGG pathway maps on the RIG-I-like receptor signalling pathway are indicated.

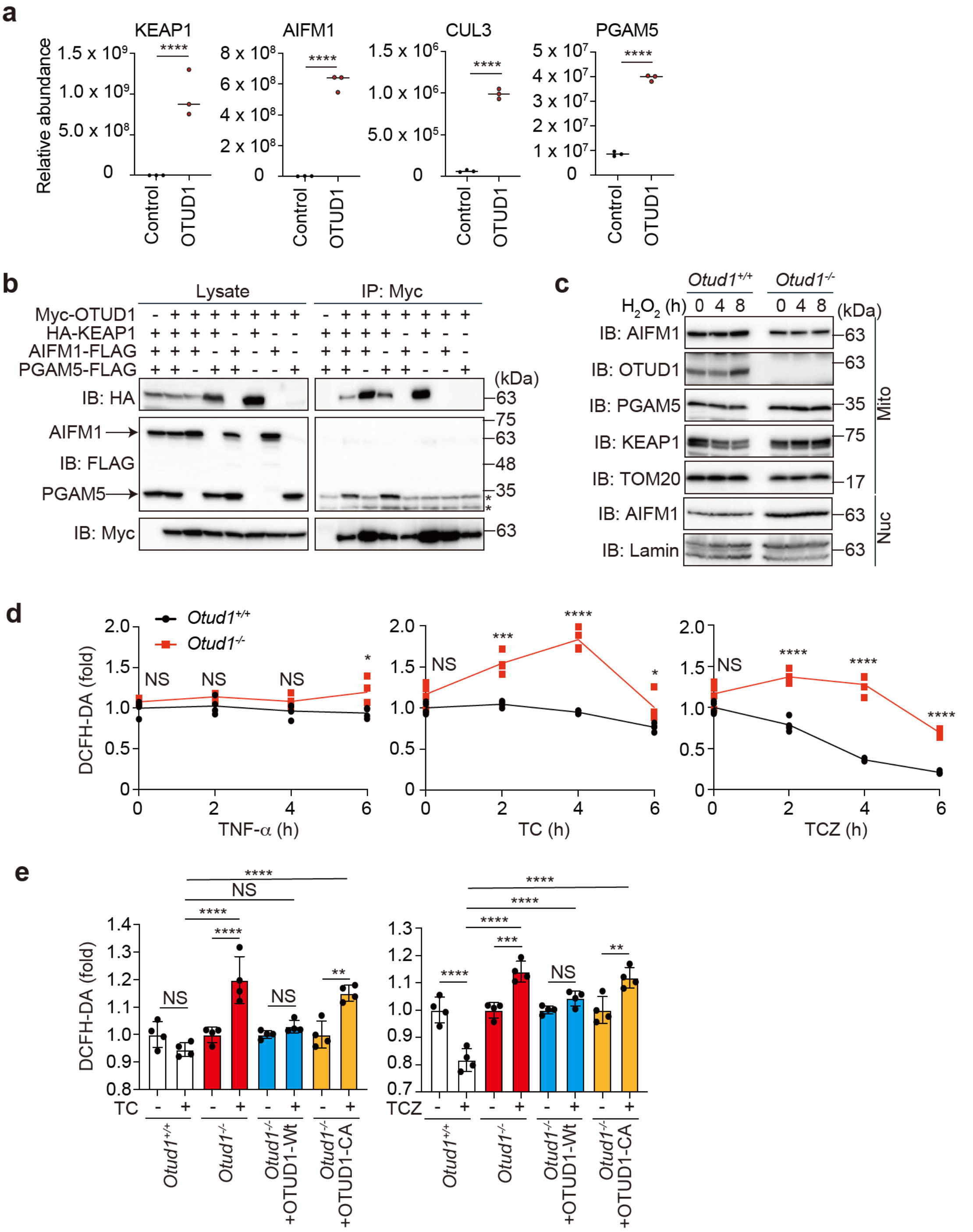


**Extended Data Fig. 5 OTUD1 regulates ROS production and cell death pathways. a**, The amounts of proteins co-precipitated with FLAG-OTUD1 were quantified by targeted mass spectrometry, using the parallel reaction monitoring (PRM) method. **b**, OTUD1 binds PGAM5 *via* KEAP1. Myc-OTUD1, HA-KEAP1, AIFM1-FLAG, and PGAM5-FLAG were expressed in HEK293T cells as indicated. The cell lysates and anti-Myc immunoprecipitates were immunoblotted with the indicated antibodies. **c**, Intranuclear localization of AIFM1 in *Otud1*$^{-/-}$-MEFs. MEFs were treated with 0.3 mM $H_2O_2$ for the indicated periods, and fractionated into mitochondrial (Mito) and nuclear (Nuc) fractions. The samples were then immunoblotted with the indicated antibodies. **d**, Enhanced ROS generation in *Otud1*$^{-/-}$ cells upon apoptotic and necroptotic stimuli. *Otud1*$^{+/+}$- and *Otud1*$^{-/-}$-MEFs were treated with 10 ng ml$^{-1}$ TNF-α, 10 μg ml$^{-1}$ CHX, and/or 20 μM ZVAD as indicated, and the intracellular ROS levels were analysed by a DCFH-DA assay. **e**, Catalytic activity of OTUD1 protects cells from ROS generation upon apoptotic and necroptotic stimuli. A similar analysis as in Fig. 5k was performed after TC or TCZ stimulation for 2 h. Data are shown as mean ± SD by *t*-test (**a**; *n* = 3, **d**; *n* = 4) or ANOVA post-hoc Tukey test (**e**; *n* = 4). *: $P$<0.05, **: $P$<0.01, ***: $P$<0.001, ****: $P$<0.0001, NS: not significant.

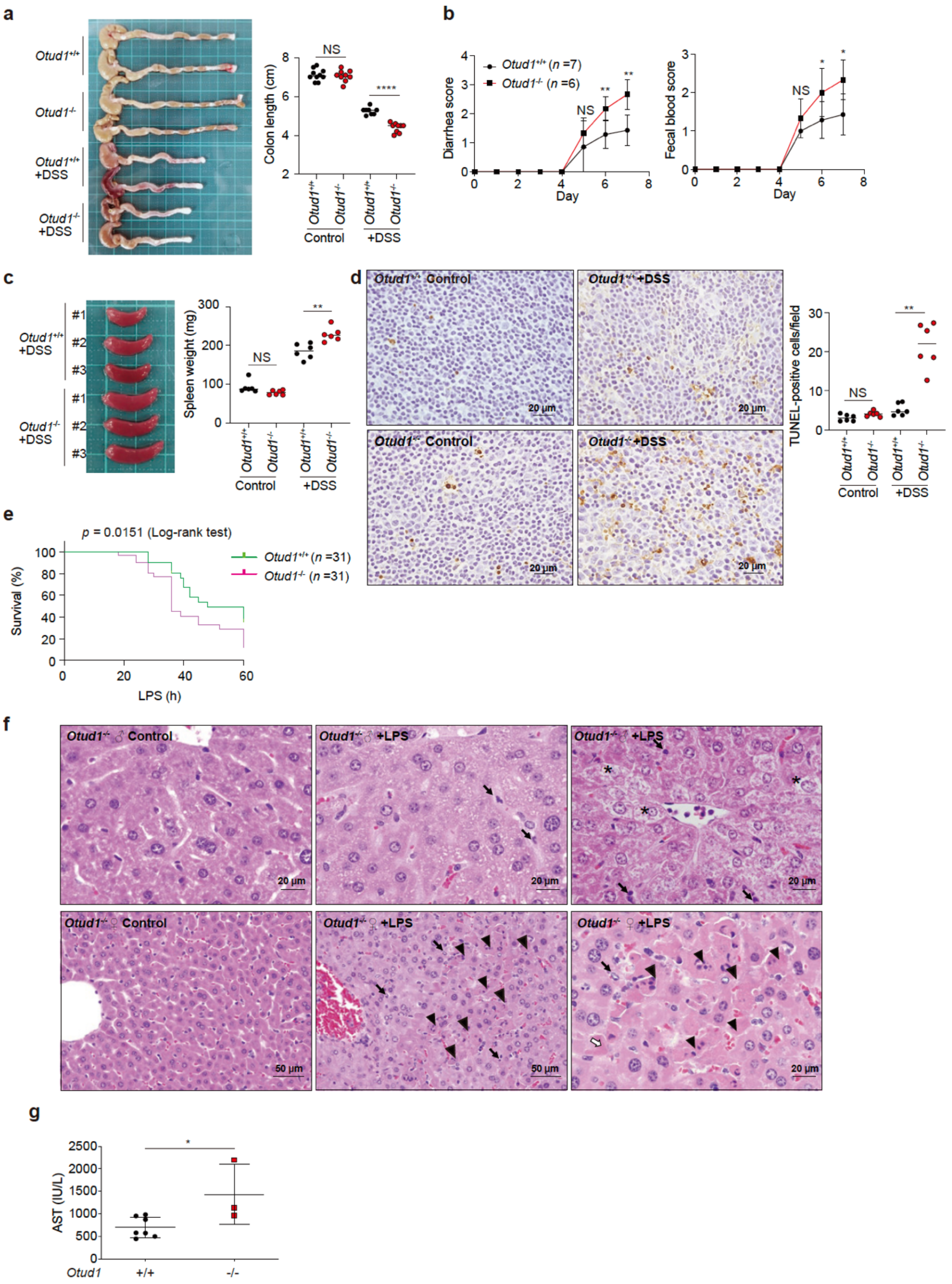

**Extended Data Fig. 6 Reduced susceptibility to pathogenic inflammatory stimuli in *Otud1*$^{-/-}$-mice. a**, Shortening of colon in DSS-administered *Otud1*$^{-/-}$-mice. Drinking water containing 2.5% DSS was administered to *Otud1*$^{+/+}$- and *Otud1*$^{-/-}$-mice for 7 days, and colon lengths were analysed. Representative images (*left*) and lengths of colons (*right*, *n* = 8-10) are shown. **b**, Increased pathology scores in DSS-treated *Otud1*$^{-/-}$-mice. The scores for diarrhea and fecal blood in *Otud1*$^{+/+}$- (*n* = 7) and *Otud1*$^{-/-}$-mice (*n* = 6) were evaluated as described in the Methods. **c**, Splenomegaly in DSS-treated *Otud1*$^{-/-}$-mice. Images (*left*) and weights of spleens (*right*, *n* = 6) are shown. **d,** Increased cell death of splenocytes in DSS-treated *Otud1*$^{-/-}$-mice. TUNEL staining of spleen sections, and numbers of TUNEL-positive cells per 40x field (*n* = 6) were analysed. *Bars*: 20 µm. **e**, *Otud1*$^{-/-}$-mice are sensitive to LPS. Survival percentages of LPS (20 mg $kg^{-1}$ i.p. injection) treated *Otud1*$^{+/+}$- and *Otud1*$^{-/-}$-mice (*n* = 31 each) determined by the Kaplan-Meier method are shown. Log-rank test was performed. **f**, Histopathological changes in the livers of *Otud1*$^{-/-}$-mice 24 h after LPS treatment. H&E stained livers from *Otud1*$^{-/-}$-male (*upper panels*) or *Otud1*$^{-/-}$-female mice (*lower panels*) with normal controls (*left panels*) or LPS-treated (*middle and right panels*) mice are shown. *Arrowheads*: necrotic hepatocytes; *arrows*: infiltration of inflammatory cells; *asterisks*: cell degradation. *Bars*: 20 or 50 µm. **g,** Increased AST level in LPS-treated *Otud1*$^{-/-}$-mice. AST levels in plasma from *Otud1*$^{+/+}$- (*n* = 7) and *Otud1*$^{-/-}$-mice (*n* = 3) after 36 h administration of LPS. **a**, **b**, **c**, **d**, **g**, Data are shown as scatter plots and evaluated by the Mann-Whitney test. *: $P<0.05$, **: $P<0.01$, ****: $P<0.0001$, NS: not significant.
